# Covalent inhibition of the SARS-CoV-2 NiRAN domain via an active-site cysteine

**DOI:** 10.1101/2024.11.22.624893

**Authors:** Genaro Hernandez, Adam Osinski, Abir Majumdar, Jennifer L. Eitson, Monika Antczak, Krzysztof Pawłowski, Hanspeter Niederstrasser, Kelly A. Servage, Bruce Posner, John W. Schoggins, Joseph A. Ready, Vincent S. Tagliabracci

**Affiliations:** Department of Molecular Biology, University of Texas Southwestern Medical Center, Dallas, Texas 75390, USA; Howard Hughes Medical Institute, University of Texas Southwestern Medical Center, Dallas, Texas 75390, USA; Department of Microbiology, University of Texas Southwestern Medical Center, Dallas, Texas 75390, USA; Department of Biochemistry, University of Texas Southwestern Medical Center, Dallas, Texas 75390, USA; Harold C. Simmons Comprehensive Cancer Center, UT Southwestern Medical Center, Dallas, Texas 75390, USA; Hamon Center for Regenerative Science and Medicine, University of Texas Southwestern Medical Center, Dallas, Texas 75390, USA

**Author notes:** Correspondence to: Vincent S. Tagliabracci, **Email:**.

**Keywords:** NiRAN, RNA capping, pseudokinase, SARS-CoV-2, COVID-19, enzyme inhibitor, inhibitor, drug screening, drug discovery, enzyme structure

## Abstract

The kinase-like NiRAN domain of nsp12 in SARS-CoV-2 catalyzes the formation of the 5’ RNA cap structure. This activity is required for viral replication, offering a new target for the development of antivirals. Here, we develop a high-throughput assay to screen for small molecule inhibitors targeting the SARS-CoV-2 NiRAN domain. We identified NCI-2, a compound with a reactive chloromethyl group that covalently binds to an active site cysteine (Cys53) in the NiRAN domain, inhibiting its activity. NCI-2 can enter cells, bind to, and inactivate ectopically expressed nsp12. A cryo-EM reconstruction of the SARS-CoV-2 replication-transcription complex (RTC) bound to NCI-2 offers a detailed structural blueprint for rational drug design. Although NCI-2 showed limited potency against SARS-CoV-2 replication in cells, our work lays the groundwork for developing more potent and selective inhibitors targeting the NiRAN domain. This approach presents a promising therapeutic strategy for effectively combating COVID-19 and potentially mitigating future coronavirus outbreaks.

## Introduction

The COVID-19 pandemic, caused by SARS-CoV-2, imposed a significant burden to our healthcare systems, societal structures, and economies (1–4). The rapid development of vaccines against SARS-CoV-2 and their subsequent updates against novel strains mitigated the number of cases of COVID-19. However, despite their high efficacy, vaccines can become obsolete if the virus evolves escape variants, and vaccine hesitancy is becoming an increasingly worrisome issue (5–8). These concerns highlight the need for comprehensive antiviral therapies to address COVID-19 and help prevent future coronavirus outbreaks.

Current antiviral drugs available for the treatment of SARS-CoV-2 target viral replication (9–14). Remdesivir and molnupiravir disrupt the SARS-CoV-2 nsp12 RNA-dependent RNA polymerase (RdRP) (9, 11, 14), while nirmatrelvir inhibits the viral nsp5 main protease (mPro), which prevents processing of the polyprotein and halts viral replication (10, 12). While these drugs have demonstrated favorable outcomes, they each have challenges and limitations, such as incomplete or partial efficacy due to viral escape variants, timing of treatment, and toxicity associated with side effects or patient comorbidities (9–11, 15–21). Therefore, it is necessary to identify new antiviral targets for the development of other classes of efficacious drugs against SARS-CoV-2 and other coronaviruses.

Coronaviruses genomes and transcripts contain a 5′ RNA cap that is essential for RNA stability, translation of viral proteins and protection against the host immune response (22, 23). To generate the RNA cap, the SARS-CoV-2 nsp12 nidovirus RdRp-associated nucleotidyl transferase (NiRAN) domain conjugates nascently transcribed RNA to nsp9 (RNAylation) and subsequently transfers the RNA to GDP, forming the core 5′cap structure GpppA-RNA (24, 25). Viral replication is abolished when mutations are introduced into key residues in the NiRAN domain or nsp9 that participate in the formation of the cap (24). Although the NiRAN domain adopts a protein kinase fold, it acts as a GDP-polyribonucleotidyltransferase that catalyzes an essential step in the viral life cycle. This unique biochemistry makes the NiRAN domain an attractive target for the development of antivirals.

The NiRAN domain can also transfer nucleotide monophosphates (NMPs) derived from nucleotide triphosphates to the N-terminus of nsp9 (NMPylation). NMPylated nsp9 may facilitate the initiation of viral RNA synthesis or protect nsp9 from degradation via the N-end rule (26–28). In the present study, we utilized the AMPylation activity of the NiRAN domain to develop a high-throughput screening assay to identify small molecule inhibitors. From this screen, we identified a compound that covalently binds to Cys53 within the NiRAN active site, inhibiting NiRAN-dependent activities.

## Results and Discussion

### Identification of NCI-1 and NCI-2 as inhibitors of the NiRAN-domain

We exploited the ability of the SARS-CoV-2 NiRAN domain to AMPylate the N-terminus of nsp9 in a high-throughput screen (HTS) to identify small molecule inhibitors of the NiRAN domain (24, 26, 27). We quantified NiRAN-dependent AMPylation of nsp9 by measuring the remaining ATP levels using the luciferase-based ATP detection reagent, Kinase-Glo Plus (**Fig. 1, *A* and *B***). We screened the entire University of Texas Southwestern Medical Center small molecule compound library, consisting of >350,000 small molecules. The screen had an average Z’ value of 0.7 (29), which is a quantitative measure of screen quality, and yielded compound hits with single-point corrected inhibition of NiRAN-dependent AMPylation scores above 10% (**Fig. 1*C***) (30). The screen yielded 24 top compound inhibitor candidates of NiRAN activity, which we used in secondary assays to test for inhibition of NiRAN-dependent nsp9 RNAylation activity (**Fig. S1**). The compounds were also assayed for binding to nsp12 via differential scanning fluorimetry, where many hits destabilized nsp12 (**Fig. S1*B-H***). Notably, however, SW157903, SW080949 (hereafter referred to as NiRAN covalent inhibitor 1; NCI-1) and SW090466 (NCI-2) showed inhibition of NiRAN-dependent RNAylation of nsp9 and an increase in the thermal stability of nsp12 (**Fig. 1*D-E* and Fig. S1*A-B, H***). By filtering out hits with pan assay interference (PAINS) substructures (31), we initially determined NCI-1 to be the only lead NiRAN inhibitor compound from our screen. However, both NCI-1 and NCI-2 are structurally similar, with both compounds featuring a fused, two-ring heterocyclic system with a chloromethyl group (**Fig. 1*F-G***). Additionally, both compounds had similar IC_50_ values against NiRAN-dependent AMPylation of nsp9 (**Fig. 1*H-I***). Therefore, we proceeded to further characterize NCI-1 and NCI-2.

**Figure 1.**
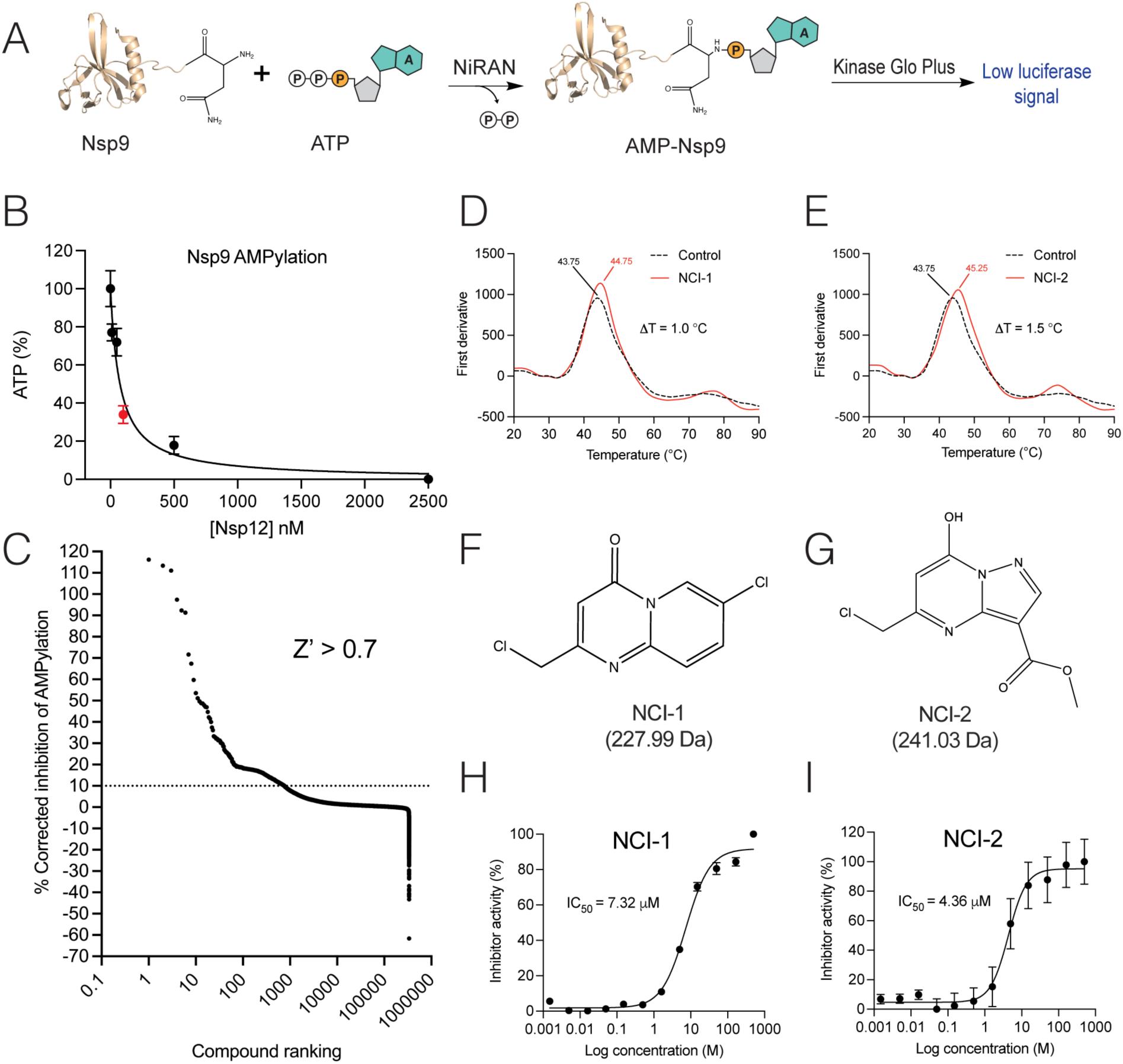
A high throughput screen (HTS) identifies inhibitors of the NiRAN domain. A, schematic representation of the catalytic reaction quantified in the HTS used in this study to identify inhibitors of NiRAN-dependent nsp9 AMPylation. B, dependence of nsp12 concentration on ATP consumption (50 µM) during NiRAN-mediated AMPylation of nsp9 (50 µM). The red point indicates the concentration of nsp12 used in the HTS. C, rank plot depicting the range of inhibition by compounds of NiRAN-catalyzed AMPylation of nsp9 in the screen. Differential scanning fluorimetry profiles of nsp12 (10 µM) in the presence of NCI-1 (20 µM) (D) and NCI-2 (20 µM) (E). Chemical structures of NCI-1 (F) and NCI-2 (G). Dose response curves of nsp12-dependent AMPylation of nsp9 with NCI-1 (H) or NCI-2 (I). The assay contained 100 nM nsp12, 50 µM nsp9 and 50 µM ATP. The reaction products were quantified via Kinase Glo Plus. The IC_50_ values are shown (n = 4). Error bars represent the standard error of the mean (SEM).

### NCI-1 and NCI-2 covalently bind Cys53 within the NiRAN active site

NCI-1 and NCI-2 contain chloromethyl functional groups predicted to react and covalently bind to the thiol group of Cys residues (32). Therefore, we hypothesized that these compounds covalently bind to nsp12. We incubated nsp12 with NCI-1or NCI-2 at a 1:1.5 protein-to-compound molar ratio and analyzed the products by intact mass spectrometry (IMS). IMS analysis revealed mass differences (Λ1m) consistent with the covalent addition of NCI-1 (**Fig. 2*A***) and NCI-2 (**Fig. 2*B***) to nsp12 involving substitution of the chloride group. Moreover, NCI-1 exhibited a second peak mass shift, suggesting the conjugation of two NCI-1 molecules to nsp12 (**Fig. 2*A***).

**Figure 2.**
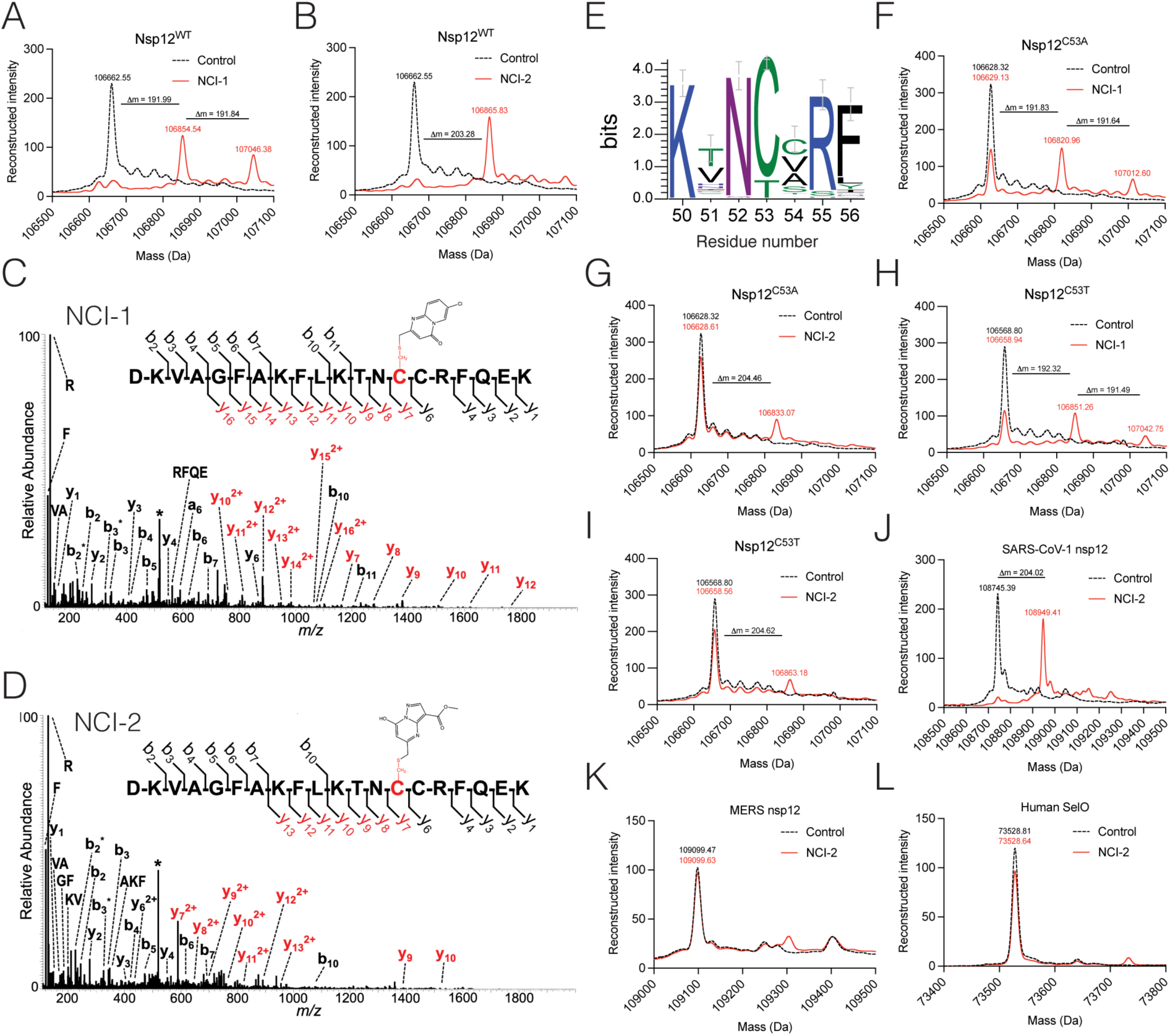
NCI-1 and NCI-2 form covalent bonds with Cys53 within the SARS-CoV-2 NiRAN domain. Intact mass LC-MS spectra of nsp12 (black trace) overlayed with nsp12 incubated with NCI-1 (red trace, A) or NCI-2 (red trace, B) at a 1:1.5 protein:drug ratio. The observed masses are shown above the respective peaks for each curve. The theoretical mass for nsp12 is 106660.24 Da. The theoretical mass shifts (Λ1m) for NCI-1 and NCI-2 bound to nsp12 are 191.53 Da and 204.57 Da, respectively. MS/MS spectra of nsp12 depicting the peptide ion DKVAGFAKFLKTNCCRFQEK with covalent attachment of either NCI-1 (+192 Da) (C) or NCI-2 (+205 Da) (D) on residue C53 of nsp12. Spectra were generated via higher-energy collisional dissociation (HCD) and fragment ions shown in red depict mass shifts corresponding to the addition of the modifying group. The precursor ion is labeled with an asterisk. Any fragment ions labeled with (*) correspond to loss of H_2_O (−18 Da) or NH_3_ (−17 Da). E, sequence logo highlighting the conservation of C53 in the NiRAN domain. The heights of the amino acid stacks correspond to the sequence conservation at given positions. Intact mass LC/MS spectra of nsp12^C53A^ (black) overlayed with nsp12^C53A^ incubated with NCI-1 (red trace, F) or NCI-2 (red trace, G). The observed masses are shown above the respective peaks for each curve. Intact mass LC/MS spectra of nsp12^C53T^ (black trace) overlayed with nsp12^C53T^ incubated with NCI-1 (red trace, H) or NCI-2 (red trace, I). The observed masses are shown above the respective peaks for each curve. Intact mass LC-MS spectra of SARS-CoV-1 nsp12 (black trace, J), MERS nsp12 (black trace, K), and human SelO^U667C^ (black trace, K) overlayed NCI-2 (red traces). The observed masses are shown above the respective peaks for each curve.

To identify the Cys residues modified by NCI-1 and NCI-2, we incubated nsp12 with the compounds and analyzed ASP-N digested peptides by LC-MS/MS. We observed several peptides corresponding to the addition of the compounds to Cys53 within the NiRAN active site (**Fig. 2*C-D***). While we detected modifications on other cysteines, the spectral counts were significantly lower compared to those for Cys53 (**Fig. S2**).

Cys53 is relatively conserved across the *Coronaviridae* family, with 80% of species containing a Cys at this position, while the remaining species have a Thr (**Fig. 2*E***). Although IMS analysis of nsp12 with Cys53 substituted to Ala or Thr revealed a mass corresponding to the unmodified protein in the presence of both compounds (**Fig. 2*F-I***), NCI-2 appeared to be more selective, with only one minor additional peak detectable in the C53 mutants (**Fig. 2*G-I***). While the SARS-CoV-2 nsp12 Cys53 equivalent is found in SARS-CoV-1 nsp12, it is not present in MERS-CoV nsp12 or human Selenoprotein-O (SelO), which shares structural similarity with the NiRAN domain (33). As expected, NCI-2 covalently binds to SARS-CoV-1 nsp12 (**Fig. 2*J***), but not to MERS nsp12 (**Fig. 2*K***), or SelO (**Fig. 2*L***). Thus, both NCI-1 and NCI-2 covalently bind to Cys53 within the NiRAN domain with NCI-2 demonstrating greater selectivity for the NiRAN active site Cys53. Consequently, we chose to concentrate on NCI-2 for the remainder of the study.

### Cys53 is required for the inhibitory activity of NCI-2

Given that NCI-2 binds to Cys53 in the NiRAN active site, we hypothesized that this residue is crucial for the compound’s ability to inhibit NiRAN-dependent activities. Substituting Cys53 with either Thr or Ala had no negative effect on NiRAN-catalyzed AMPylation of nsp9 (**Fig. 3*A***), RNAylation of nsp9 (**Fig. 3, *B* and *C***) or core cap formation, as judged by the deRNAylation of nsp9 (**Fig. 3*D***) (24). However, the inhibitory effect of NCI-2 on all three of these activities was abolished in both Cys53 mutants (**Fig. 3, *A, C,* and *D***). Notably, NCI-2 did not impact nsp12 polymerase activity (**Fig. 3*E***). Thus, while Cys53 is dispensable for NiRAN activity, it is essential for the inhibitory effect of NCI-2.

**Figure 3.**
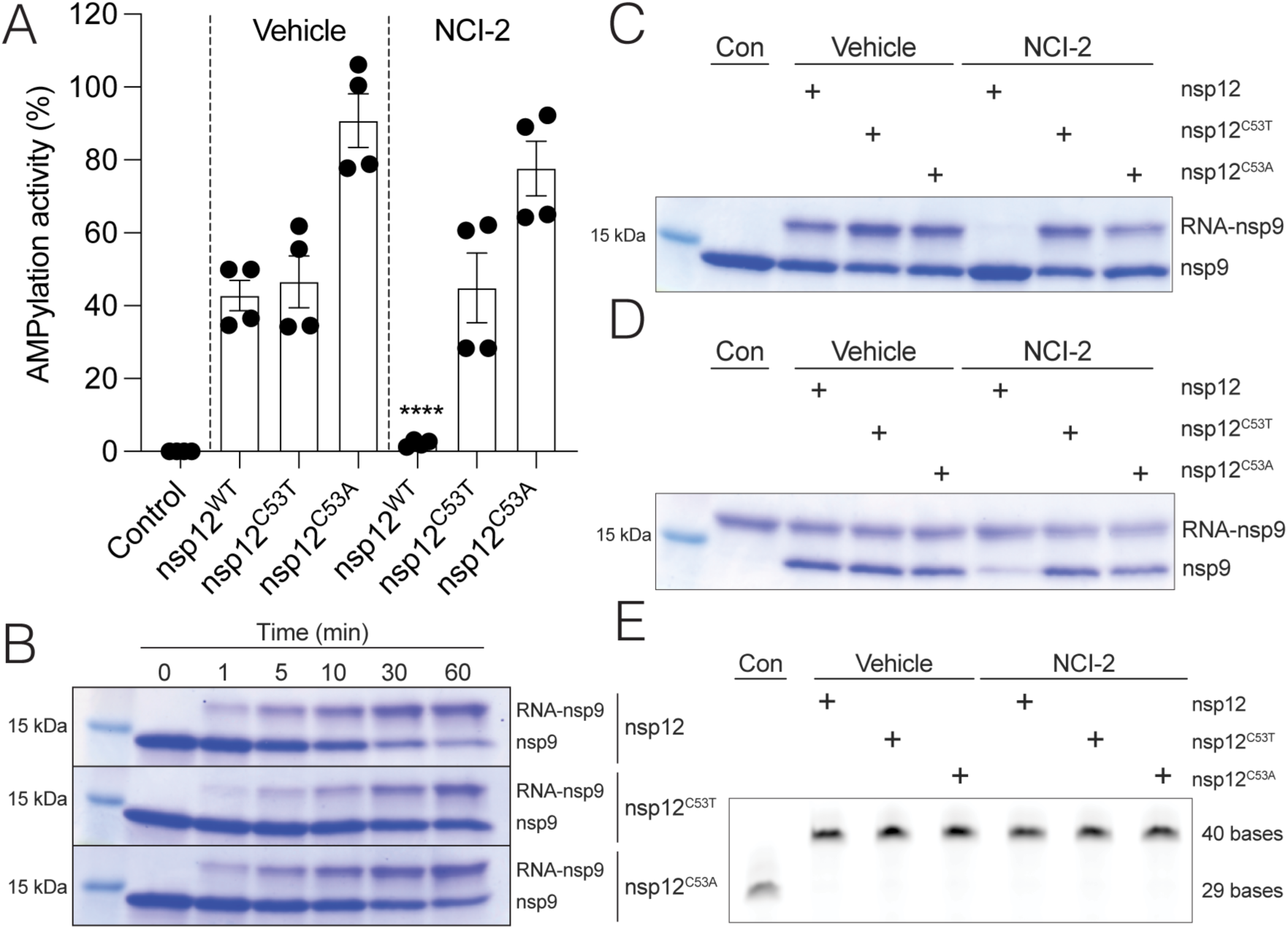
Cys53 within the NiRAN domain is essential for the inhibitory activity of NCI-2. A, AMPylation activity of nsp12, nsp12^C53T^ or nsp12^C53A^ (100 nM) in the presence of NCI-2 (20 μM). Reactions were allowed to proceed for 30 minutes, and the products were analyzed by Kinase Glo Plus (n = 4; error bars indicate SEM; **** p < 0.0001 for the indicated group compared against all other groups except the no enzyme control group). B, Time-dependent RNAylation of nsp9 (20 μM) by nsp12, nsp12^C53T^ or nsp12^C53A^ (500 nM). Reactions were initiated with 100 μM of SARS-CoV-2 genomic leader sequence corresponding to the first 10 bases in the genome, and the reaction products were analyzed by SDS-PAGE and Coomassie staining. C, RNAylation activity of nsp12, nsp12^C53T^ or nsp12^C53A^ (500 nM) following incubation with NCI-2 (20 μM) for 30 min prior to the start of the reaction. Reaction products were analyzed as in (B). D, nsp12, nsp12^C53T^ or nsp12^C53A^ (500 nM)-mediated deRNAylation of nsp9 (20 μM) following incubation with NCI-2 (20 μM) for 30 min prior to the start of the reaction. Reaction products were analyzed as in (B). Note that the deRNAylation of nsp9 is a readout of core cap GpppA formation (24). E, nsp12, nsp12^C53T^ or nsp12^C53A^ (500 nM) RdRp activity in an RNA extension activity. nsp12 and mutants were preincubated with the cofactors nsp7 (5 μM), nsp8 (5 μM), NCI-2 (20 μM) for 30 min, initiated by the addition of a fluorescently labelled self-priming hairpin RNA and NTPs. The reaction products were separated by UREA-PAGE and analyzed by fluorimetry.

### NCI-2 demonstrates limited potency in inhibiting SARS-CoV-2 replication

To assess whether NCI-2 can inhibit nsp12 in cells, we first confirmed that it is non-toxic to HEK293A cells at concentrations up to 200 µM after 8 hours of treatment (**Fig. 4*A***). LC-MS/MS analysis confirmed the cellular uptake of NCI-2 (**Fig. 4*B***). Therefore, we tested whether NCI-2 can bind nsp12 in cells by synthesizing the NCI-2 analog SW395943 (NCI-3) that contains a click-reactive alkyne handle attached to the ester functional group of NCI-2 (**Fig. 4*C***). Similar to NCI-2, NCI-3 has an IC_50_ of 10.41 µM for NiRAN-dependent AMPylation of nsp9 and displayed no cellular toxicity (**Fig. 4, *D* and *E***). We treated HEK293A cells stably expressing an N-terminally tagged nsp12 (Flag-nsp12) with NCI-3, conjugated a biotin to the alkyne group of NCI-3 in cell lysates and analyzed protein biotinylation by means of SDS-PAGE and avidin-HRP. While we observed a major biotinylated species corresponding to nsp12, we also observed nonspecific binding of NCI-3 to other proteins (**Fig. 4*F***). To determine if NCI-2 inhibits NiRAN activity in cells, we immunoprecipitated FLAG-nsp12 from cells treated with NCI-2 and evaluated RNAylation activity using nsp9 as the substrate. Indeed, Flag-nsp12 immunoprecipitated from NCI-2 treated cells, showed a marked reduction in its ability to RNAylate nsp9, indicating that NCI-2 effectively inhibited FLAG-nsp12’s activity in cells. (**Fig. 4*G***).

**Figure 4.**
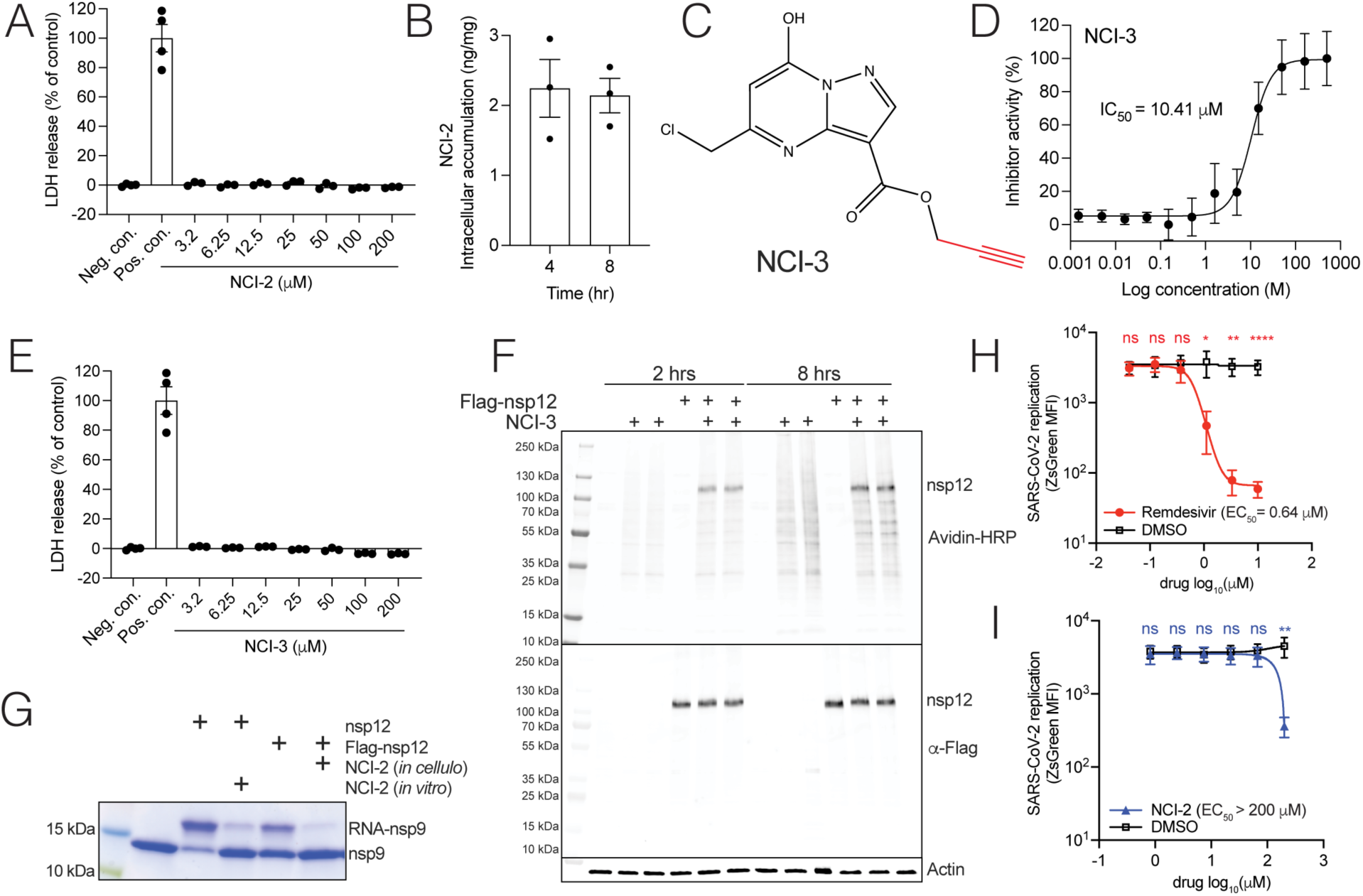
NCI-2 enters cells, inhibits ectopically expressed nsp12, and shows limited potency against SARS-CoV-2. A, LDH release assay as a readout of cell toxicity following treatment of HEK293A cells with NCI-2 for 8 hours at 37°C and 5% CO_2_, with vehicle (1% DMSO) treated cells as a negative control and lysed cells as positive control. B, intracellular accumulation of NCI-2 in cells at 4 and 8 hours post treatment. HEK293A cells cultured in serum free medium, were incubated with NCI-2 (80 μM) for 30 minutes and NCI-2 was quantified by LC-MS/MS analysis. C, chemical structure of the NCI-2 derivative NCI-3 depicting the alkyne handle (red). D, dose response curve of nsp12-dependent AMPylation of nsp9 with NCI-3. Reactions were analyzed as in Fig. 1H. The IC_50_ value is shown. E, LDH release assay as a readout of cell toxicity following treatment of HEK293A cells with NCI-3 (80 μM). F, analysis of HEK293A cell lysates for biotinylation via avidin-HRP or nsp12 via protein immunoblotting following treatment with NCI-3 (80 μM). Following 2- and 8-hour incubations, cells were harvested, and the lysates underwent a copper-catalyzed click reaction with biotin azide to generate biotin-NCI-3 conjugates for detection. Actin is shown as a loading control. G, RNAylation activity of Flag-tagged nsp12 (Flag-nsp12) immunoprecipitated from HEK293A lysates following treatment of the cells with 80 μM NCI-2 (*in cellulo*). Purified nsp12 (500 nM) was also included in these assays as a control and treated with NCI-2 (80 μM) for 30 minutes prior to the start of the reaction (*in vitro*). Reaction products were analyzed as in Fig. 3B. Infectivity of SARS-CoV-2-Wu-1-zsGreen in the presence of varying concentrations of remdesivir (H) or NCI-2 (I). A549-TMPRSS2-ACE2 cells were infected with SARS-CoV-2 for 30 min at 37°C. Cells were washed to remove virus and medium containing DMSO and the indicated drugs were added back. At 7 hours post-infection, cells were fixed with 4% PFA. Viral infectivity was quantified by flow cytometry. N = 3 biological replicates. Statistical analysis was determined by paired t-test. *, P<0.05; **, P<0.01; ****, P<0.0001, ns, not significant.

To assess whether NCI-2 can inhibit SARS-CoV-2 replication, we infected A549 cells expressing ACE2 and TMPRSS2 with SARS-CoV-2 carrying a ZsGreen reporter. We then treated the infected cells with either NCI-2 or remdesivir at 30 minutes post-infection and measured ZsGreen levels as a readout of viral replication by flow cytometry. While remdesivir inhibited viral replication with an EC_50_ of 0.64 μM (**Fig. 4*H***), we only observed inhibition of viral replication by NCI-2 at 200 μM (**Fig. 4*I***). Thus, although NCI-2 can enter cells, bind to, and inactivate ectopically expressed nsp12, it exhibits low potency against SARS-CoV-2 replication in cells.

### The inhibitory activity of NCI-2 is dependent on its chloromethyl group

We conducted a dose-dependent time course of NCI-2 inhibition of nsp12 NiRAN-dependent AMPylation, determined the rate of inactivation constant (*k*_inact_ = 0.1441 min^-1^), inhibition constant (K_i_ = 16.32 μM) and second order rate constant (*k*_inact_ / K_i_ = 8.83 ×10^-3^ min^-1^ μM^-1^) (**Fig 5, *A* and *B***). These data suggest that NCI-2 is a low potency, covalent inhibitor with moderate-to-weak covalent reactivity (*k*_inact_ < 3 min^-1^) and low affinity for its target (K_i_ > 50 nM for 100 nM nsp12 in the assay reaction) (34). Structure-activity IC_50_ analyses of NCI-2 derivatives with the Cys-reactive chloromethyl group substituted to either a less reactive or non-reactive functional group confirmed that the chloromethyl group reactivity is essential for NCI-2 inhibition of NiRAN activity (**Fig. 5*C***). We next explored whether substitutions of the non-reactive hydroxyl and methyl ester functional groups of NCI-2 would help increase its target affinity and enhance the compound’s inhibitory capacity. NCI-2 hydroxyl and methyl ester group substitutions led to a decrease (IC_50_ > 500 μM) or complete loss (no inhibitory activity up to 500 μM) of inhibitory capacity (**Fig. 5, *D* and *E***). Thus, NCI-2 is a weak covalent inhibitor that requires its Cys-reactive chloromethyl group to covalently inhibit NiRAN activity.

### Structural insights into the binding of NCI-2 to the SARS-CoV-2 NiRAN domain

We determined a cryo-EM structure of the SARS-CoV-2 replication/transcription complex (RTC) bound to NCI-2 (**Fig. 6 and Fig. S3, S4, Table 1**). The Coulomb potential map clearly shows the density for the ligand and the covalent bond with Cys53 (**Fig. 6*A*)** NCI-2 occupies the active site space responsible for interactions with nucleotide substrates and sterically prevents the “base-in” mode of binding (**Fig. 6, *B* and *C***) (35). While there are no significant clashes with the “base-up” nucleotide orientation (**Fig. 6*C***), several residues typically involved in catalysis and/or nucleotide phosphate binding adopt different orientations (Lys50, Lys73, Arg116) (**Fig. 6, *D* and *E***).

**Figure 5.**
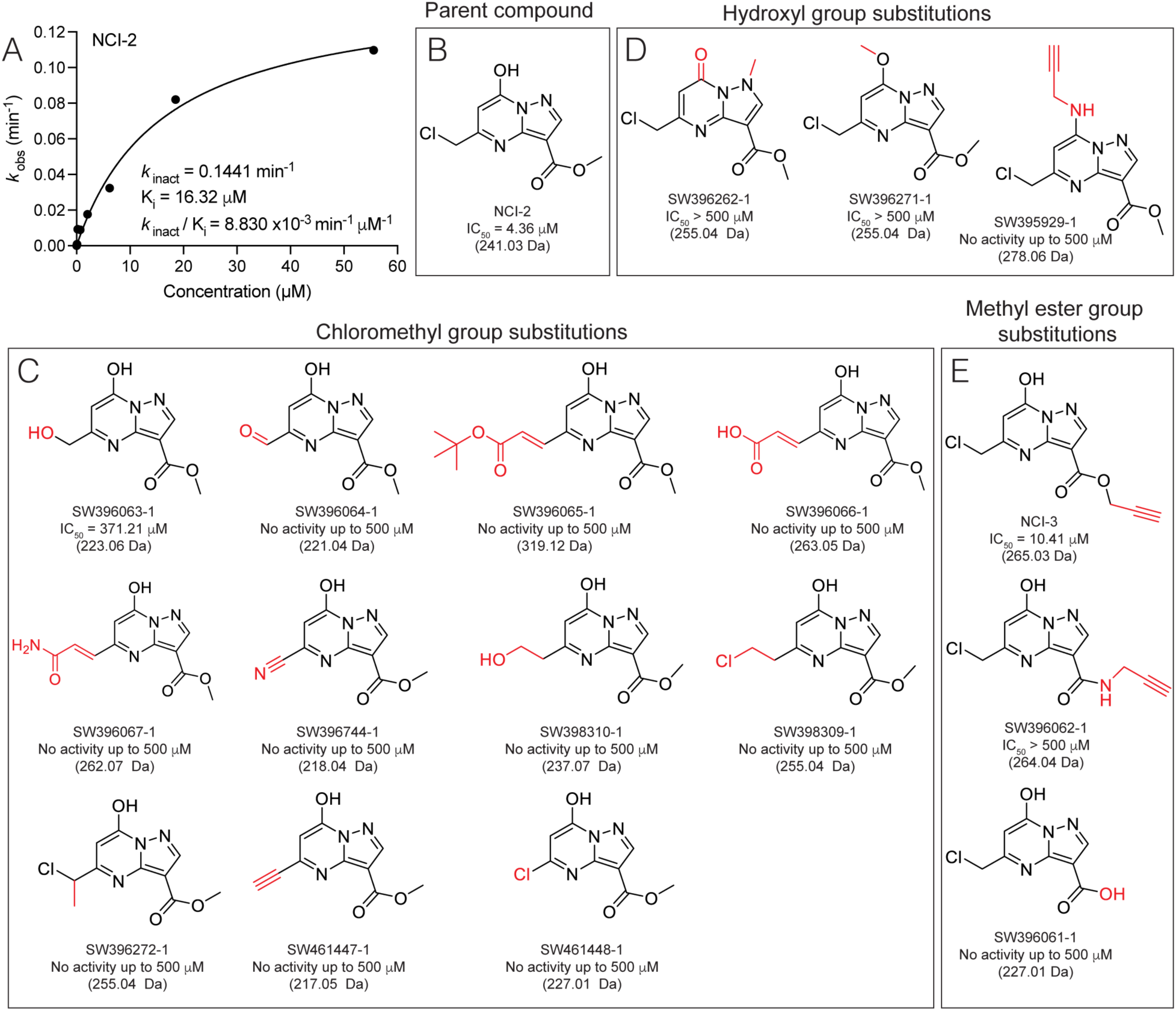
Structure-activity analysis of NCI-2. A, kinetic analysis curve of a time and dose-dependent assay with calculated values for *k*_inact_, K_i_, and *k*_inact_ / K_i_ for NiRAN-dependent nsp9 AMPylation inhibition activity with 100 nM nsp12 measured via Kinase Glo Plus (n = 4). B, chemical structure of NCI-2 with indicated IC_50_ values for inhibition of nsp9 AMPylation. NCI-2 derivatives with the chloromethyl (C) hydroxyl (D) or methyl ester (E) functional groups substituted to the functional group indicated in red for each structure. IC_50_ values quantified as in Fig. 1I are shown.

**Figure 6.**
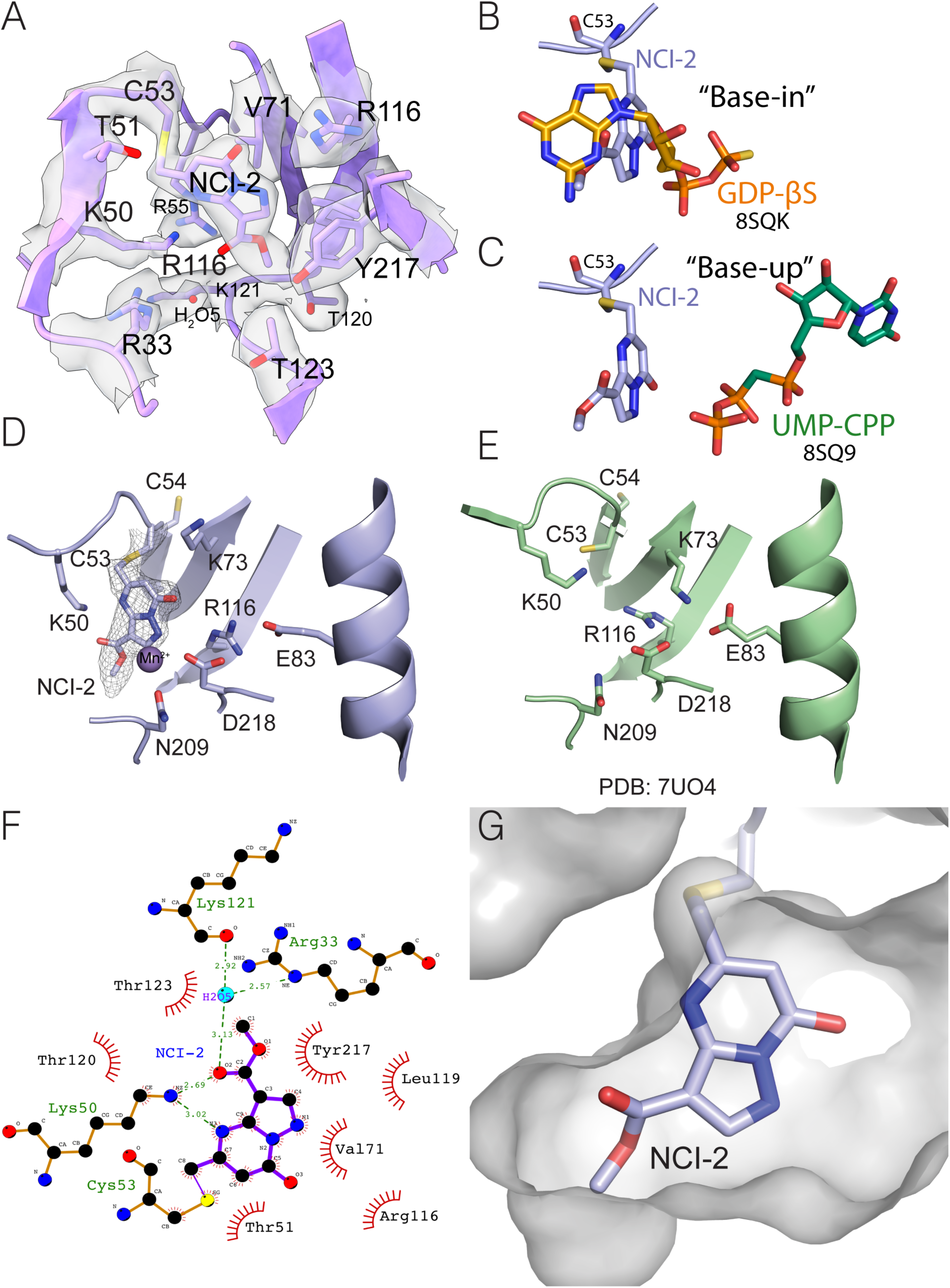
Cryo-EM structure of NCI-2 bound to Cys53 in the NiRAN domain. A, cartoon representation of the NiRAN domain active site bound to the covalent inhibitor NCI-2 with the coulombic density presented as a transparent surface. Superimposition of nucleotides bound in the NiRAN active site in “base-in” (B) and “base-up” (C) conformations (PDBID 8SQ9). Cartoon representations of bound NCI-2 (D) and an apo structure of the NiRAN domain (E) (PDB: 7UO4) (36). Select residues important for catalysis are shown as sticks. Coulombic density for the compound and the Cys53 is shown as mesh. Map was sharpened with a B-factor = −71.2. F, LigPlot (60) showing interactions between NCI-2 and the NiRAN domain. G, surface representation of the NiRAN active site proximal to the inhibitor, shown as sticks.

**Table 1.** Cryo-EM data collection, refinement and validation statistics.

|  |  |
| --- | --- |
|  | RTC + SW090466-1<br>(EMDB-45587)<br>(PDB 9CGV) |
| **Data collection and processing** |  |
| Magnification | 105kx |
| Voltage (kV) | 300 |
| Electron exposure (e-/Å<sup>2</sup>) | 60 |
| Defocus range (µm) | 1.1-2.4 |
| Pixel size (Å) | 0.84 |
| Symmetry imposed | C1 |
| Initial particle images (no.) | 8M |
| Final particle images (no.) | 77,971 |
| Map resolution (Å) | 2.7 |
| FSC threshold | 0.143 |
| Map resolution range (Å) | 2.7-7 |
| **Refinement** |  |
| Initial model used (PDB code) | 7UO4 |
| Model resolution (Å) | 2.7 |
| FSC threshold | 0.143 |
| Model resolution range (Å) | 2.7-4.8 |
| Map sharpening *B* factor (Å<sup>2</sup>) | -71.2 |
| Model composition |  |
| Non-hydrogen atoms | 10404 |
| Protein residues | 1179 |
| Ligands | 1 |
| *B* factors (Å<sup>2</sup>) |  |
| Protein | 47.57 |
| Ligand | 67.03 |
| R.m.s. deviations |  |
| Bond lengths (Å) | 0.004 |
| Bond angles (°) | 0.474 |
| Validation |  |
| MolProbity score | 0.91 |
| Clashscore | 1.62 |
| Poor rotamers (%) | 0 |
| Ramachandran plot |  |
| Favored (%) | 98.2 |
| Allowed (%) | 1.8 |
| Disallowed (%) | 0 |

NCI-2 binding causes major rearrangements of the active site when compared to the apo structure (**Fig. 6, *D*, and *E*, Fig. S4, *G* and *H***). The catalytic Lys73 is displaced from the salt bridge with Glu83 and is replaced by Arg116 that normally interacts with the nucleotide phosphates (36). Lys50 moves towards the back of the active and forms electrostatic interactions with NCI-2 (**Fig. 6*F***). Additional electrostatic interactions are formed through an ordered water molecule (H_2_O5) with the Arg33 guanidinium and a main chain carbonyl of Lys121. The heterocyclic ring of the inhibitor forms extensive interactions with a hydrophobic patch formed by Val71 and Leu119, while the methyl ester group forms hydrophobic interactions with Tyr217 and Thr123 (**Fig. 6*F***). Binding-induced rearrangements also resulted in displacement and destabilization of a strand neighboring Cys53, which becomes poorly ordered (missing residues 23-29 with no visible density, **Fig. S4, *G* and *H***).The overall changes upon NCI-2 binding likely result in a large energy penalty during the initial binding event, contributing to the poor binding and kinetics of the compound. Moreover, there are no strong electrostatic interactions that facilitate NCI-2 binding to the active site. Additional available space was observed for derivative development at the hydroxyl and methyl ester functional group positions of NCI-2 (**Fig. 6*G***). Collectively, our structural analysis of NCI-2 binding to the NiRAN domain provides a preliminary blueprint for advancing compound optimization.

## Conclusion

In this work, we developed a biochemical HTS assay that allowed us to screen for small molecule inhibitors of NiRAN-dependent nsp9 AMPylation and identified NCI-2 as a covalent inhibitor of the NiRAN domain. NCI-2 has a two-ring heterocyclic structure that resembles purines and a chloromethyl reactive group that binds Cys53 within the NiRAN active site. While NCI-2 could enter cells, engage ectopically expressed nsp12 and inhibit NiRAN activity, it displayed limited potency against SARS-CoV-2 in cells. Thus, further optimization will be needed to improve NCI-2. Nevertheless, the identification of NCI-2, its mechanism of action, and the structural elucidation of its binding to the NiRAN domain represent a promising foundation that should catalyze future drug development efforts targeting the NiRAN domain. Furthermore, we anticipate that NCI-2 can be used as a tool compound to study RNA capping by SARS-CoV-2.

## Experimental procedures

### Chemicals and reagents

Anti-Flag M2 Affinity Gel (A2220), Flag M2 monoclonal antibody (F3165), 3X Flag peptide (F4799), Chloramphenicol (C0378), kanamycin sulfate (K1377), Glycerol (G7757), Antifoam B emulsion (A5757), dithiothreitol (DTT; D0632), imidazole (I2399), IPTG (I5502), 2-mercaptoethanol (BME, M3148), Brilliant Blue R (B0149), manganese (II) chloride tetrahydrate (MnCl_2_; M3634), magnesium chloride (MgCl_2_; M2670), potassium chloride (KCl; P9541), Asp-N endoproteinase (11420488001), SYPRO Orange, toxoflavin (K4394), and urea (U6504), were obtained from Millipore-Sigma. Q5 DNA polymerase (M0492L), all restriction enzymes used for cloning, proteinase K (P8107S), yeast inorganic pyrophosphatase (M2403) were obtained from New England Biolabs. ATP, acetic acid (A38-212), 2× TBE-urea sample buffer (LC6876), HisPur Ni-NTA resin (88223) were obtained from Thermo Fisher Scientific. Phenylmethylsulfonyl fluoride (PMSF; 97064-898) was from VWR. 4–20% Mini-PROTEAN TGX Stain-Free Protein Gels (4568096) were obtained from BioRad. SW080949 (NCI-1; K093-0001) and SW090466 (NCI-2; K784-2867) were obtained from ChemDiv.

### Plasmids and bacmids

SARS-CoV-2 nsp7, nsp8, nsp12, SARS-CoV-1 nsp12, and MERS nsp12 coding sequences (CDSs) were codon-optimized for bacterial, insect, mammalian cell expression, and synthesized as gBlocks (Integrative DNA Technologies). The CDS for nsp9 was amplified from a mammalian expression vector (a gift from N. Krogan) (37). The CDSs were cloned into modified pET28a bacterial expression vectors containing N-terminal 6/8×His tags followed by the yeast Sumo (smt3) CDS. Amino acid mutations were introduced using QuikChange site-directed mutagenesis. In brief, primers were designed using the Agilent QuikChange primer design program to generate the desired mutation and were used in PCR reactions with PfuTurbo DNA polymerase. The reaction products were digested with Dpn1, transformed into DH5α cells and mutations were confirmed by Sanger sequencing.

For protein expression in *Escherichia coli*, ppSumo-SARS-CoV-2 nsps and mutants were cloned into a BamH1 site at the 5′ end, which introduced a Ser residue following the diGly motif in smt3. To make native N termini, the codon encoding the Ser was deleted using QuickChange mutagenesis. Thus, after cleavage with the ULP protease (after the diGly motif), the proteins contained native N termini.

For expression in insect cells the nsp12 CDSs for SARS-CoV-2, SARS-CoV-1, and MERS were subcloned into a pFastBac Dual vector (Thermo Fisher Scientific) with a C-terminal 10×His tag. The final sequence was confirmed by Sanger sequencing. Cloned nsp12 pFastBac Dual vectors were transformed into DH10Bac competent cells (Thermo Fisher Scientific) to generate bacmids for bacuolovirus production.

For stable cell protein expression in mammalian cells the SARS-CoV-2 CDS was cloned into a pQCXIP retroviral expression vector with an N-terminal FLAG tag. The final sequence was verified by Sanger sequencing.

### Protein expression and purification in *Escherichia coli*

#### nsp7, nsp8, nsp12, and nsp12 Cys53 mutants

SARS-CoV-2 6×His-Sumo-nsp7/8, 8×His-Sumo-nsp12 and the corresponding mutant plasmids (with native N termini following the diGly motif in Sumo) were transformed into Rosetta (DE3) *E. coli* cells under 50 μg/ml kanamycin exposure. LB Miller growth medium starter cultures (5 ml) containing 50 μg/ml kanamycin and 34 μg/ml chloramphenicol were grown for 2–4 h at 37 °C and then transferred to growth medium containing the same antibiotics. Protein expression was induced at an optical density (OD) of 0.7–1.0 by adding 0.4 mM IPTG and overnight incubation (16 h) at 18 °C. Cultures were centrifuged at 3,000 x g for 10 min and the bacterial pellet was resuspended in lysis buffer (50 mM Tris pH 8.0, 300 mM NaCl, 1 mM PMSF, 15 mM imidazole pH 8.0 and 1 mM DTT) and lysed by sonication. Lysates were centrifuged for 30 minutes at 30,000 x g and the supernatants incubated with Ni-NTA resin in a nutator for 1 h at 4 °C. The Ni-NTA resin was washed with 50 mM Tris, pH 8.0, 300 mM NaCl, 30 mM imidazole pH 8.0, 1 mM DTT and the protein was eluted in 50 mM Tris pH 8.0, 300 mM NaCl, 300 mM imidazole pH 8.0 and 1 mM DTT. The eluted proteins were incubated with 5 μg/ml Ulp1 protease overnight at 4 °C. nsp7 was separated from 6×His–Sumo by anion exchange (Capto HiRes Q 5/50 column (Cytiva) equilibrated in 50 mM Tris pH 8.0, 50 mM NaCl, 1 mM DTT, eluted with 0–50% gradient of buffer containing 1 M NaCl). 6×His–SUMO–nsp8 was treated with Ulp1 on the Ni-NTA resin, to separate nsp8 and 6×His–SUMO in buffer with no imidazole. Proteins were further purified by size-exclusion chromatography using a Superdex 200 10/300 increase, Superdex 200 16/600, Superdex 75 10/300 increase or Superdex 75 16/600 column (all columns from Cytiva) in 50 mM Tris pH 8.0, 150 mM NaCl and 1 mM DTT. The fractions containing proteins of interest were pooled, concentrated in an Amicon Utra-15 centrifugal filters (pore size, 3–50 kDa). Purified proteins were aliquoted and stored at −80 °C.

### Purification of nsp9

6×His-Sumo-nsp9 (with native N termini following the diGly motif in Sumo) was transformed into Rosetta (DE3) *E. coli* cells under 50 μg/ml kanamycin exposure. LB Miller growth medium starter cultures (5 ml) containing 50 μg/ml kanamycin, and 34 μg/ml chloramphenicol were grown for 2–4 h at 37 °C and then transferred to Terrific Broth (TB) growth medium containing the same antibiotics and several drops of Antifoam B emulsion. Protein expression was induced at an OD of 1.2 by adding 0.4 mM IPTG and incubating overnight for 16 hours at 18 °C. The cultures were centrifuged at 3,000 x g for 10 min and the bacterial pellet was resuspended in 50 mM Tris, pH 8.0, 300 mM NaCl, 5% glycerol, 1 mM PMSF, 15 mM imidazole pH 8.0, 1 mM DTT, and lysed by sonication. Lysates were centrifuged for 30 minutes at 30,000 x g and the supernatants incubated in Ni-NTA resin for 1 hour at 4 °C. The Ni-NTA resin was washed with 50 mM Tris, pH 8.0, 300 mM NaCl, 30 mM imidazole pH 8.0 and 1 mM DTT, followed by a low salt wash in 50 mM Tris pH 8.0, 60 mM NaCl, 30 mM imidazole pH 8.0 and 1 mM DTT. The protein was eluted in 50 mM Tris pH 8.0, 60 mM NaCl, 300 mM imidazole pH 8.0 and 1 mM DTT and incubated with 5 μg/ml Ulp1 overnight at 4 °C. The protein was diluted with 50 mM Tris pH 8.0, 1 mM DTT buffer to lower the NaCl concentration to 30 mM and subsequently ran through a Hi-Trap CaptoQ (Cytiva) column where the flowthrough contained purified nsp9. NaCl was added to each protein to a final concentration of 150 mM, concentrated in an Amicon Ultra-15 with a 10 kDa molecular weight cut-off (MWCO), aliquoted and stored at −80 °C.

### Purification of RNA-nsp9 species

RNA-nsp9 was purified as previously described (24). Briefly, purified native nsp9 (0.8 mg/mL, 65 μM) was incubated at room temperature overnight with 130 μM of 5’ triphosphorylated SARS-CoV-2 genomic leader sequence 10-mer RNA and ∼0.8 μM of nsp12 in presence of 0.05 mg/ml yeast inorganic pyrophosphatase and 1 mM MnCl_2_, in the reaction buffer (50 mM Tris pH 7.5, 5 mM KCl, 1 mM DTT). The samples were clarified by centrifugation to remove any precipitate and applied directly onto a Capto HiRes Q 5/50 column (Cytiva) equilibrated in 50 mM Tris pH 8.0, 50 mM NaCl and 1 mM DTT. An elution gradient of 0–50% with 1 M NaCl was applied over 30 column volumes. Under these conditions, RNA and RNA-nsp9 bound to the column and unmodified nsp9 did not. RNA-nsp9 and unreacted RNA eluted as a peak doublet around 70 mS cm−1. The fractions were pooled, and further purified over Superdex 75 increase 10/300 GL (50 mM Tris pH 8.0, 300 mM NaCl, 1 mM DTT), separating RNA-nsp9 from unreacted RNA and nsp12. RNA-nsp9 was quantified by spectrophotometry with an estimated extinction coefficient of ɛ260 = 130,650 M−1 cm−1.

### Baculovirus production, insect cell protein expression and purification

SARS-CoV-2, SARS-CoV-1 or MERS bacmids were transfected into adherent Sf21 cells (Expression Systems) using Cellfectin II transfection reagent at a 1:1 bacmid:Cellfectin II ratio and incubated in Grace’s unsupplemented medium (Thermo Fisher Scientific) for 5 hours at 27 °C. The medium was then replaced with Grace’s supplemented medium (Thermo Fisher Scientific) with 10% FBS and the cells were kept at 27°C for 48 hours. The medium containing P1 baculovirus was transferred to Sf21 cells in suspension in ESF921 medium (Expression Systems) to generate P2 virus in an orbital shaker at 110 rpm, 27°C for 72 hours. 5 ml of P2 virus were transferred to a fresh 500 ml culture of suspension Sf21 cells to generate P3 virus at 110 rpm, 27°C for 72 hours.

For protein expression, 10 ml of P3 baculovirus were added to 1 L of Sf21 suspension cells in ESF921 at 110 rpm, 27°C for 72 hours. Cells were centrifuged at 2000 x g for 10 minutes and resuspended in 50 mM Tris pH 8.0, 300 mM NaCl, 5% glycerol, 1 mM DTT, 1 mM PMSF, 15 mM imidazole pH 8.0, and 1× Complete protease inhibitor cocktail (Roche). Cells were lysed via sonication and the lysate cleared by centrifugation at 30,000 x g for 30 minutes. The lysates were then incubated with Ni-NTA resin in a nutator for 1 hour at 4°C. The resin was washed twice with 50 mM Tris pH 8.0, 300 mM NaCl, 1 mM DTT, and 30 mM imidazole pH 8.0 followed by protein elution in 50 mM Tris pH 8.0, 300 mM NaCl, 1 mM DTT, 300 mM imidazole pH 8.0, and concentrated in an Amicon Ultra-15 centrifugal filter with a 50 kDa MWCO. Proteins were further purified by size-exclusion chromatography using a Superdex 200 10/300 increase column or Superdex 200 16/600 in 50 mM Tris pH 8.0, 150 mM NaCl and 1 mM DTT. The fractions containing nsp12 were pooled and concentrated in an Amicon Ultra-15 centrifugal filter with a 50 kDa MWCO, aliquoted and stored at −80 °C.

### Generation of stable Flag-nsp12 HEK293A cells

HEK293A cells (ATCC) were maintained in complete medium (DMEM supplemented with 10% FBS and 1% penicillin/streptomycin). Cells were co-transfected with pQCXIP-Flag-nsp12 and pCL-10A1 retroviral packaging vector with PolyJet (SignaGen) transfection reagent at a 3:1 DNA:PolyJet ratio and incubated for 5 hours at 37°C in a CO_2_-controlled cell incubator. Cell medium was replaced with fresh complete medium, and cells were allowed to recover for 2 days to allow for retrovirus production at 37°C. The medium containing retrovirus was then transferred to newly plated HEK293A cells and incubated for 2 days at 37°C. Cells stably expressing Flag-nsp12 were isolated by puromycin (5 μg/ml) selection on complete medium and protein expression was confirmed by immunoblotting.

### LDH release assay

HEK293A cells were plated in 96-well plates at a density of 10,000 cells per well in DMEM. The cells were treated with a 2-fold dilution series of NCI-2 or NCI-3 ranging from 200 µM to 3.125 µM and DMSO only control for 8 hours at 37°C. 100 µl of medium from each were then transferred to a new 96-well plate and 100 μl of CytoTox 96 colorimetric reagent (Promega) were added to each well and incubated for 30 minutes at room temperature in the dark. LDH release was measured by the absorbance at 490 nm in a Spark Cyto microplate reader. The negative control for baseline used was medium devoid of cells. The positive control for 100% LDH release was generated by scrapping and lysing the cells via passage through a 26 ½ gauge syringe needle prior to medium collection.

### AMPylation assay for high-throughput compound screening and drug kinetic analysis

For the small molecule high-throughput compound screen of NiRAN-dependent nsp9 AMPylation inhibitors, the UT Southwestern Medical Center compound library comprising more than 350,000 small, drug-like compounds was screened at the institutions High Throughput Screening Core Facility. Compounds were dry spotted in 384-well white bottom plates (Greiner Bio-One) using an Echo 655 acoustic liquid handler (Beckman, Inc.). Then 5 μl of 2× protein reaction mix (50 mM Tris pH 7.5, 5 mM KCl, 1 mM DTT, 100 µM nsp9 and 200 nM nsp12) was dispensed into each well of the plates using an automated dispenser (BioTek MultiFlo liquid handler, Agilent, Inc.) and incubated for 30 minutes at room temperature. The reactions were started by the addition of 5 μl of a 2× reaction start mix (50 mM Tris pH 7.5, 5 mM KCl, 1 mM DTT, 100 μM ATP and 2 mM MnCl_2_) using the same automated dispenser and incubated for 30 minutes at room temperature. The final reaction volume per well was 10 μl with each compound having a final concentration of 5 μM with the vehicle DMSO being 0.1%, 5 µM toxoflavin as a positive control and buffer only negative control (one replicate per compound). The final concentration in the reaction for nsp9 and ATP was 50 μM and 100 nM for nsp12 (except for when varying concentrations of nsp12 were tested). 10 μl of Kinase Glo Plus (Promega) were then added using the same automated dispenser to stop the reactions and incubated for 30 minutes at room temperature. The luciferase signal was measured in a CLARIOstar plus microplate reader (BMG Labtech) and the data analyzed using Genedata Screener (Genedata) software. For all assay plates in the screen, the Z’ values were greater than 0.7 (average Z’ value across all plates was 0.82). Compound hits were defined as having >10% inhibition of AMPylation and Z-score < −3. Hit validation was conducted under the same protocol as the primary screen with the top ∼500 compounds tested at 15, 5, and 1.25 μM with three (3) replicates per dose. Compounds that failed to show dose-responsiveness or inhibitory activity were filtered out.

AMPylation assays for the calculation of drug IC_50_ values for all compounds and when nsp12 C53 mutants were used in this study were conducted under the same protocol as the HTS assay. For kinetic and inhibition constant determination of NCI-2, varying drug preincubation times were performed with a concomitant dose-response, with no automated volume dispensing and luciferase signal measured in a Spark Cyto microplate reader (Tecan), with all other conditions being the same as the HTS assay. Graphpad Prism software (Graphpad) was used to calculate IC_50_, *k*_inact_, K_i_, and *k*_inact_ / K_i_ (38).

### Differential scanning fluorimetry

10 μM of nsp12 protein was mixed with 20 μM of the indicated inhibitor or DMSO in a buffer containing 50 mM Tris pH 7.5, 5 mM KCl, 1 mM DTT, 1 mM MnCl_2_ and 20× SYPRO Orange. Samples were run in triplicate in a CFX Opus 384 real-time PCR system (BioRad) under a melting curve program with a 0.5°C/second ramp from 20°C to 90°C. The raw RFU values were then analyzed in Graphpad Prism to calculate the first derivative and identify the melting curve peak temperatures.

### RNAylation assay

RNAylation reactions were carried out in a 10 µl volume containing 50 mM Tris pH 7.5, 5 mM KCl, 1 mM DTT, 2.5 U/ml yeast inorganic pyrophosphatase, 20 µM nsp9 and either 500 nM nsp12 or Cys53 nsp12 mutants. The proteins were preincubated with either vehicle DMSO, 20 μM NCI-2, NCI-1 or the other top hit compounds from the HTS for 30 minutes at room temperature. The reactions were started by adding MnCl_2_ and 5’ triphosphorylated SARS-CoV-2 genomic leader sequence 10-mer RNA (5’-pppATTAAAGGTT-3’; ChemGenes) to a final concentration of 100 µM. The reactions were incubated for 30 minutes at 37 °C and terminated with the addition of 5× SDS-PAGE sample buffer + BME and boiling the samples for 5 minutes. The reaction products were resolved by SDS–PAGE on a 4–20% gradient gel and visualized by Coomassie staining.

### Intact-mass analysis

Samples were prepared in a 20 μl final volume containing 50 mM Tris pH 7.5, 5 mM KCl, 1 mM DTT, 1 mM MnCl_2_, either 10 μM SARS-CoV-2 nsp12, SARS-CoV-2 nsp12 Cys53 mutants, SARS-CoV-1 nsp12, MERS nsp12 or human SelO were incubated with 15 μM of NCI-1 or NCI-2 for at least 1 hour at room temperature. The samples were then analyzed at the UT Southwestern Proteomics core facility by LC–MS, using a Sciex X500B Q-TOF mass spectrometer coupled to an Agilent 1290 Infinity II HPLC. Samples were injected onto a POROS R1 reverse-phase column (2.1 mm × 30 mm, 20 µm particle size, 4,000 Å pore size) and desalted. The mobile phase flow rate was 300 μl min−1 and the gradient was as follows: 0–3 min, 0% B; 3–4 min, 0–15% B; 4–16 min, 15–55% B; 16–16.1 min, 55–80% B; 16.1–18 min, 80% B. The column was then re-equilibrated at the initial conditions before the subsequent injection. Buffer A contained 0.1% formic acid in water and buffer B contained 0.1% formic acid in acetonitrile.

The mass spectrometer was controlled by Sciex OS v.1.6.1 using the following settings: ion source gas 1, 30 psi; ion source gas 2, 30 psi; curtain gas, 35; CAD gas, 7; temperature, 300 °C; spray voltage, 5,500 V; declustering potential, 80 V; collision energy, 10 V. Data were acquired from 400–2,000 Da with a 0.5 s accumulation time and 4 time bins summed. The acquired mass spectra for the proteins of interest were deconvoluted using BioPharmaView v.3.0.1 (Sciex) to obtain the molecular mass values. The peak threshold was set to ≥5%, reconstruction processing was set to 20 iterations with a signal-to-noise threshold of ≥ 20 and a resolution of 2,500. The final reconstructed protein mass traces were plotted using Graphpad prism.

### Identification of cysteine residues in nsp12 modified by NCI-1 and NCI-2 by mass spectrometry

Samples were prepared in a 20 μl final volume containing 50 mM Tris pH 7.5, 5 mM KCl, 1 mM DTT, 1 mM MnCl_2_, and 10 μM SARS-CoV-2 nsp12 were incubated with 20 μM of NCI-1 or NCI-2 for at least 1 hour at room temperature. Protein samples were resolved by SDS–PAGE on a 4–20% gradient gel, visualized by Coomassie staining, and excised for mass spectrometry analysis. Proteins were reduced with 10 mM DTT for 1 hour at 56°C and then alkylated with 50 mM iodoacetamide for 45 minutes at room temperature in the dark. Overnight enzymatic digestion at 37°C was performed with Asp-N. Peptide samples were de-salted via solid phase extraction (SPE) prior to analysis by LC-MS/MS. Experiments were performed on a Thermo Scientific EASY-nLC liquid chromatography system equipped with a 50 cm Thermo PepMap analytical column (2 µm particle size, 75 µm diameter) (Thermo Fisher Scientific) and a Orbitrap Fusion Lumos mass spectrometer (Thermo Fisher Scientific). MS1 spectra were acquired in the Orbitrap mass analyzer (resolution 120,000) and peptide precursor ions were isolated and fragmented using high-energy collision-induced dissociation (HCD) to produce MS2 spectra. MS/MS fragmentation spectra were acquired in the ion trap. Spectral data was searched using the Mascot search engine (Matrix Science) against the sequence for SARS-CoV-2 nsp12 protein. The precursor mass tolerance was 15 ppm and the fragment mass tolerance was 0.6 Da. Variable modifications included carbamidomethylation of cysteine (+57.021 Da) and oxidation of methionine (+15.995 Da). Additionally, two new modifications were added and searched for covalent inhibition modification of cysteine by NCI-1 inhibitor (+192.009 Da) and NCI-2 inhibitor (+205.049 Da). The identification of all peptides modified by either inhibitor were manually verified to confirm the correct assignment of modification site.

### Intracellular concentration measurement of NCI-2 by mass spectrometry

HEK293A cells were incubated in DMEM with 20 µM NCI-2. At 4 or 8 hours post compound addition, media was removed, and cells were washed twice with cold PBS; cells were trypsinized, rinsed with 2 ml cold medium and transferred to a conical tube on ice. Cells were spun down and resuspended in cold PBS. They were counted and spun down a second time and resuspended at 1×10^6^ cells/ml. Cells were snap frozen and transferred to the PPC. 100 µL of lysate was mixed with 200 µL of methanol containing 0.15% formic acid and 75 ng/mL tolbutamide. Tubes were vortexed, incubated at room temperature for 10 min, and spun twice at 16,100 x g. Supernatant was analyzed by LC-MS/MS at the UT Southwestern Preclinical Pharmacology core facility. The protein pellet after addition of methanol was saved, dried, resuspended in 0.1N NaOH and subjected to a BCA assay (Pierce) for protein content using a BSA standard curve. Data were normalized to cell pellet protein content.

### NCI-3 detection in HEK293 cells by western blot

Wildtype HEK293A cells or cells stably expressing Flag-nsp12 were treated with 80 μM NCI-3 or DMSO control in DMEM for 2 or 8 hours. Cells were washed twice with PBS and lysed in RIPA buffer + protease inhibitor cocktail. Cell lysates were cleared by centrifugation at 21,000 x g for 15 minutes. Total protein concentration was measured in the cleared lysates with a DC protein assay kit (BioRad) and the concentrations were adjusted with RIPA buffer to normalize all sample concentrations. Click chemistry reactions to detect NCI-3 were performed for all samples. Briefly, 20 µl reactions containing 3 mg/ml cleared lysate, 100 µM TBTA, 2 mM CuSO_4_, 1 mM sodium ascorbate, 25 µM biotin azide, RIPA buffer + protease inhibitor cocktail, were incubated for 1 hour in the dark. The reaction products were then resolved by SDS–PAGE on a 4–20% gradient gel and transferred to a nitrocellulose membrane (BioRad). The membrane was blocked with 2.5 % BSA in TBST for 1 hour at room temperature with gentle rocking. Antibodies against Flag (raised in mouse) and Actin (raised in rabbit) (Cell Signaling Techonolgy) were incubated with the membrane overnight at 4°C with gentle rocking. The following day, the membrane was washed 3 times with TBST for 5 minutes each time, then incubated with Alexa Fluor 488 anti-rabbit (Thermo Fisher Scientific), anti-mouse IRDye 800CW (LI-COR Biosciences) secondary antibodies, as well as with IRDye 680RD streptavidin (LI-COR Biosciences) for 1 hour at room temperature in the dark with gentle rocking. The membrane was washed 3 times with TBST for 5 minutes each time. The fluorescent signal from the membrane corresponding to Flag, actin and streptavidin were imaged in an Oddyssey M system (LI-COR Biosciences).

### Flag-nsp12 immunoprecipitation

HEK293A cells were treated with 80 μM NCI-2 or DMSO control in DMEM for 8 hours. Cells were then washed twice with PBS and harvested in 25 mM Tris pH 8.5, 140 mM NaCl, 0.1% Tween 20, and lysed via passage through a 26-1/2 gauge syringe. The lysates were cleared by centrifugation at 21,000 x g for 15 minutes at 4°C. 30 µl of anti-Flag M2 affinity gel was added to each cleared lysate and incubated for 2 hours at 4°C in a nutator. The Flag M2 affinity gel was then washed 4 times with TBS and the protein was eluted with 100 µg/ml 3X Flag peptide for 1 hour at 4°C in a nutator. The eluted protein was concentrated in a 0.5 ml Amicon Ultra-0.5 centrifugal filter unit with Ultracel-30 membrane. Purified protein concentrations were estimated by absorbance at 260 nm and 0.01 mg/ml from each sample was used for the RNAylation assay as described above. Insect cell purified recombinant nsp12 (500 nM) pretreated with 20 µM NCI-2 for 30 minutes at room temperature was used as a positive control.

### Coronavirus NiRAN sequence logo generation

Sequence logo for the Cys53 region of the SARS-CoV-2 NiRAN domain was created using the WebLogo method (39). Sequences of SARS-CoV-2 nsp12 homologs were collected in the NCBI RefSeq database by BLAST search and made non-redundant by clustering with mmseqs (40) at 0.9 sequence identity level, aligned by Mafft (41) and edited by removing alignment columns containing gaps in the query sequence.

### DeRNAylation assay

DeRNAylation reactions were performed in a 10 µl reaction volume consisting of 50 mM Tris pH 7.5, 5 mM KCl, 1 mM DTT, 20 µM RNA-nsp9, and 500 nM wildtype or Cys53 nsp12 mutants. The reactions were preincubated with either vehicle DMSO or 20 µM of the indicated small molecule compound for 30 minutes at room temperature. The reactions were then started by adding 1 mM MgCl_2_, 500 µM GDP and incubated at 37 °C for 30 minutes. The reactions were stopped by addition of 5× SDS–PAGE sample buffer + BME and boiling the samples for 5 minutes. The reaction products were resolved by SDS– PAGE on a 4–20% gradient gel and visualized by Coomassie staining.

### RNA extension assay

Nsp12 RdRP-dependent RNA extension reactions were performed as previously described with some modifications (42). Reactions were set up in a 10 µl volume consisting of 50 mM Tris pH 8.0, 5 mM KCl, 1 mM DTT, 0.2 U/µl RNAse inhibitor, 5 µM nsp7, 5 µM nsp8, and 500 nM wildtype or Cys53 nsp12 mutants. The reactions were preincubated with either vehicle DMSO or 20 µM of the indicated small molecule compound for 30 minutes at room temperature. In parallel, a 5’ fluorescently labeled self-priming hairpin RNA (5’-6-FAM-UUUUCAUGCUACGCGUAGUUUUCUACGCG-3’) was annealed by heating to 75 °C and gradually cooling to 4 °C. The reactions were then started with the addition of 5 µM self-priming hairpin RNA, 1 mM MgCl_2_, 400 µM ATP, 200 µM GTP, 200 µM CTP, 200 µM UTP and incubated at 37°C for 30 minutes. The reactions were then stopped by adding 1ul of proteinase K and incubated at 37°C for 30 minutes, 2× TBE-urea Sample Buffer was then added to the samples followed by boiling for 5 minutes. The reaction products were resolved in a 15% UREA-PAGE gel and the 6-FAM signal from the RNA visualized in a ChemiDoc gel imager (BioRad).

### SARS-CoV-2 infection assays

A single cell clone (8D6) that is highly permissive to SARS-CoV-2 was generated from A549-TMPRSS2-ACE2 cells (a gift from C. Rice). Cells were maintained in DMEM supplemented with 10% FBS, 1X non-essential amino acids (NEAA), and 0.4 mg/ml Geneticin (43).

Virus from infectious clone pCC1-4K-SARS-CoV-2-Wuhan-Hu-1-ZsGreen (a gift from S. Wilson) was generated and propagated as described (44, 45). SARS-CoV-2 infections were performed in a Biosafety Level 3 (BSL3) facility equipped with the appropriate safety features to prevent biohazardous exposure or unintentional pathogen release. Experiments and biosafety protocols were reviewed and approved by the Institutional Biosafety Committee, according to guidelines provided by the UT Southwestern Office of Business and Safety Continuity.

A 3-fold dilution series of NCI-2 ranging from 200 µM to 0.82 µM, was prepared in DMEM containing 1X NEAA. Additionally, a 3-fold dilution series of Remdesivir, ranging from 10 µM to 0.04 µM, was used as a positive inhibition control. DMSO was used as a negative control. A549-TMPRSS2-ACE2-8D6 cells were plated at a density of 100,000 cells/well. Cells were infected with 1.5 MOI SARS-COV-2-zsGreen for 30 minutes at 37 °C in DMEM containing 1X NEAA. Cells were washed three times with PBS and 500 μl DMEM containing 1X NEAA and DMSO or drug dilutions was added back to cells. Cells were incubated at 37°C for 7 hours, detached with Accumax, and fixed in a final concentration of 4% PFA. Samples were analyzed by flow cytometry and viral replication was quantified as the average mean fluorescence intensity (MFI) of the ZsGreen signal in the single cell population.

### Synthesis of NCI-2 derivatives

#### NCI-3 (SW395943). Prop-2-yn-1-yl 5-(chloromethyl)-7-hydroxypyrazolo[1,5-a]pyrimidine-3-carboxylate

Prop-2-yn-1-yl 3-amino-1H-pyrazole-4-carboxylate (230 mg, 1.39 mmol) and methyl 4-chloro-3-oxobutanoate (315 mg, 2.09 mmol) were stirred at 80 °C in AcOH for 4 hours. The reaction mixture was cooled down to room temperature and filtered. The crude product was purified on a pre-packed Silica gel column on a Teledyne ISCO purification system to give desired product in 5% isolated yield. ^1^H NMR (600 MHz, DMSO) δ 12.43 (s, 1H), 8.29 (s, 1H), 6.23 (s, 1H), 4.95 (d, *J* = 2.5 Hz, 2H), 4.83 (s, 2H), 3.62 (t, *J* = 2.4 Hz, 1H). ESI-MS (m/z): 266.1 [M+H]^+^

#### SW395929. Methyl 5-(chloromethyl)-7-(prop-2-yn-1-ylamino)pyrazolo[1,5-a]pyrimidine-3-carboxylate

To the solution of methyl 7-chloro-5-(chloromethyl)pyrazolo[1,5-a]pyrimidine-3-carboxylate (24 mg, 92 μmol) in 2 ml of *i*PrOH was added propargylamine at room temperature. The reaction mixture was stirred at room temperature for 1h, followed by 20 minutes at 65 °C and 30 min at 100 °C. The reaction mixture was then filtered and solid was washed with small amount of MeOH to give 10 mg of product. ^1^H NMR (600 MHz, DMSO) δ 8.92 (t, *J* = 6.1 Hz, 1H), 8.57 (s, 1H), 6.65 (s, 1H), 4.77 (s, 2H), 4.28 (dd, *J* = 6.0, 2.5 Hz, 2H), 4.06 (s, 1H), 3.79 (s, 3H). ^13^C NMR (151 MHz, DMSO) δ 162.81, 160.27, 147.95, 147.65, 147.28, 100.68, 88.36, 79.63, 75.30, 51.39, 47.06, 31.47). ESI-MS (m/z): 279.1 [M+H]^+^

#### SW396061. 5-(Chloromethyl)-7-hydroxypyrazolo[1,5-a]pyrimidine-3-carboxylic acid. 3-amino-1H-pyrazole-4-carboxylic acid

(200 mg, 1.57 mmol) and methyl 4-chloro-3-oxobutanoate (355 mg, 2.36 mmol) were stirred at 80 °C in 3 ml of AcOH for 4 hours. The reaction mixture was cooled down to room temperature and filtered to give desired product as a solid in 78 % isolated yield. ^1^H NMR (400 MHz, DMSO) δ 12.93 (s, 1H), 12.20 (s, 1H), 8.21 (s, 1H), 6.17 (s, 1H), 4.84 (s, 2H). ESI-MS (m/z): 228.0 [M+H]^+^

#### SW396062. 5-(Chloromethyl)-7-hydroxy-N-(prop-2-yn-1-yl)pyrazolo[1,5-a]pyrimidine-3-carboxamide

To the solution of 5-(chloromethyl)-7-hydroxypyrazolo[1,5-a]pyrimidine-3-carboxylic acid (100 mg, 0.44 mmol) in 5.0 ml of DCM was added 34 μl of DMF followed by 113 μl of oxalyl dichloride at 0° C. After 30 minutes of stirring at room temperature the reaction mixture was evaporated to give 5-(chloromethyl)-7-hydroxypyrazolo[1,5-a]pyrimidine-3-carbonyl chloride.

5-(chloromethyl)-7-hydroxypyrazolo[1,5-a]pyrimidine-3-carbonyl chloride was then dissolved in 2 ml of DCM and prop-2-yn-1-amine (2.0 mmol, 110 mg,130 μl) was added. The reaction mixture was stirred for 30 minutes at 60 °C, evaporated and purified on a pre-packed Silica gel column on a Teledyne ISCO purification system. ^1^H NMR (600 MHz, DMSO) δ 8.59 (t, *J* = 5.7 Hz, 1H), 7.98 (s, 1H), 5.71 (s, 1H), 4.53 (s, 2H), 4.12 (dd, *J* = 5.7, 2.5 Hz, 2H), 3.13 (t, *J* = 2.5 Hz, 1H). ESI-MS (m/z): 265.1 [M+H]^+^

#### SW396063. Methyl 7-hydroxy-5-(hydroxymethyl)pyrazolo[1,5-a]pyrimidine-3-carboxylate

Methyl 5-((benzyloxy)methyl)-7-hydroxypyrazolo[1,5-a]pyrimidine-3-carboxylate was stirred in EtOAc at 60 °C until it was completely dissolved. The solution was then cooled down to room temperature and Pd/C (10%) was then added. The hydrogenation reaction was performed for 48h under 80-100 psi. ^1^H NMR (600 MHz, DMSO) δ 11.62 (s, 1H), 8.21 (s, 1H), 5.97 (s, 1H), 5.82 (s, 1H), 4.54 (d, *J* = 5.2 Hz, 2H), 3.83 (s, 3H. ^13^C NMR (151 MHz, DMSO) δ 162.68, 156.46, 156.13, 144.08, 143.53, 96.86, 95.71, 59.61, 51.73.). ESI-MS (m/z): 224.1 [M+H]^+^

#### SW396064. Methyl 5-formyl-7-hydroxypyrazolo[1,5-a]pyrimidine-3-carboxylate

To the solution of Methyl 7-hydroxy-5-(hydroxymethyl)pyrazolo[1,5-a]pyrimidine-3-carboxylate (24 mg, 0.11 mmol) in 2 ml of DCM was added DMP (68 mg, 0.16 mmol, 1.5 equiv.) at 0°. The reaction mixture was stirred at room temperature for 1 hour, then diluted with DCM and washed with saturated NaHCO_3_, followed by washing with saturated Na_2_O_2_S_3_. The organic layer was dried over Mg_2_SO_4_, filtered, and concentrated under reduce pressure. The resulting residue was purified on a pre-packed Silica gel column on a Teledyne ISCO purification system followed by reverse phase Teledyne ISCO purification system to provide desired product in 23 % isolated yield. ^1^H NMR (400 MHz, MeOD) δ 8.13 (s, 1H), 6.06 (s, 1H), 5.51 (s, 1H), 3.82 (s, 3H). ESI-MS (m/z): 220.1 [M-H]^-^

#### SW396065. Methyl (*E*)-5-(3-(tert-butoxy)-3-oxoprop-1-en-1-yl)-7-hydroxypyrazolo[1,5-a]pyrimidine-3-carboxylate

To the solution of methyl 5-formyl-7-hydroxypyrazolo[1,5-a]pyrimidine-3-carboxylate (110 mg, 0.50 mmol) in DCM was added tert-butyl 2-(triphenyl-l5-phosphaneylidene)acetate (225 mg, 0.60 mmol) and the reaction mixture was stirred at room temperature for 2 hours. The reaction was then diluted with DCM and washed with water. The organic layer was dried over Mg_2_SO_4_, filtered, and concentrated under reduce pressure. The resulting residue was purified on a pre-packed Silica gel column on a Teledyne ISCO purification system to give desired product in 30% isolated yield. ^1^H NMR (600 MHz, MeOD) δ 8.27 (m, 1H), 7.64 (d, *J* = 15.9 Hz, 1H), 6.82 (d, *J* = 15.8 Hz, 1H), 6.42 (s, 1H), 3.93 (s, 3H), 1.57 (s, 9H). ESI-MS (m/z): 320.2 [M+H]^+^

#### SW396066. (*E*)-3-(7-hydroxy-3-(methoxycarbonyl)pyrazolo[1,5-a]pyrimidin-5-yl)acrylic acid

To solution of methyl (*E*)-5-(3-(tert-butoxy)-3-oxoprop-1-en-1-yl)-7-hydroxypyrazolo[1,5-a]pyrimidine-3-carboxylate (30 mg, 94 μmol) in 2 ml of DCM was added TFA (180 μmol)and the reaction mixture was stirred at room temperature for 1 hour. The reaction mixture was then evaporated under reduce pressure to give desire product as a TFA salt. ^1^H NMR (600 MHz, DMSO) δ 12.05 (s, 1H), 8.28 (s, 1H), 7.83 (d, *J* = 16.0 Hz, 1H), 6.91 (d, *J* = 16.0 Hz, 1H), 6.60 (s, 1H), 3.85 (s, 3H). ^13^C NMR (151 MHz, DMSO) δ 166.84, 162.31, 155.87, 146.59, 144.04, 143.33, 135.60, 128.56, 97.73, 97.72, 51.81, 40.39, 40.25, 40.11, 39.97, 39.83, 39.69, 39.55. ESI-MS (m/z): 262.0 [M-H]^-^

#### SW396067. Methyl (*E*)-5-(3-amino-3-oxoprop-1-en-1-yl)-7-hydroxypyrazolo[1,5-a]pyrimidine-3-carboxylate

To a solution of (*E*)-3-(7-hydroxy-3-(methoxycarbonyl)pyrazolo[1,5-a]pyrimidin-5-yl)acrylic acid (10 mg, 38 μmol) in DCM was added 2.9 μl of DMF followed by 9.8 μl oxalyl chloride at 0°C. The reaction was stirred at room temperature for 30 minutes. The reaction was then quenched by the addition of NH_3_ solution in dioxide and the reaction mixture was stirred at room temperature overnight. The reaction was then evaporated and purified on a pre-packed Silica gel column on a Teledyne ISCO purification system to give desire product in 20% isolated yield. ^1^H NMR (600 MHz, DMSO) δ 8.04 (s, 1H), 7.74 (s, 1H), 7.17 (d, *J* = 15.4 Hz, 1H), 7.13 (s, 1H), 6.92 (d, *J* = 15.4 Hz, 1H), 5.81 (s, 1H), 3.71 (s, 3H). ESI-MS (m/z): 263.0 [M+H]^+^

#### SW396262. Methyl 5-(chloromethyl)-1-methyl-7-oxo-1,7-dihydropyrazolo[1,5-a]pyrimidine-3-carboxylate

To the solution of Methyl 5-(chloromethyl)-7-hydroxypyrazolo[1,5-a]pyrimidine-3-carboxylate (100 mg, 0.41 mmol) in MeOH (0.5 mL) and DCM (2.5 mL) and added 2.0 M TMS-diazomethane in hexanes (70.9 mg, 0.62 mmol, 310 ml) over 1 minute and the reaction was stirred for 20 minutes at room temperature and 1 hour at 50 °C. The reaction was then quenched with acetic acid and concentrated in vacuo to give a crude product which was purified on a pre-packed Silica gel column on a Teledyne ISCO purification system to give desired product in 5 % isolated yield. ^1^H NMR (400 MHz, CDCl_3_) δ 8.14 (s, 1H), 6.39 (s, 1H), 4.56 (s, 2H), 4.50 (s, 3H), 3.96 (s, 3H). ESI-MS (m/z): 256.0 [M+H]^+^

#### SW396271. Methyl 5-(chloromethyl)-7-methoxypyrazolo[1,5-a]pyrimidine-3-carboxylate

To the solution of methyl 5-(chloromethyl)-7-hydroxypyrazolo[1,5-a]pyrimidine-3-carboxylate (0.83 mmol, 200 mg) in 1.0 ml of POCl_3_ was added N,N-dimethylaniline (0.83 mmol, 105μl) and the reaction mixture was stirred at 100 °C for 1 hour. POCl_3_ was then evaporated and the crude mixture was purified on on a pre-packed Silica gel column on a Teledyne ISCO to give methyl 7-chloro-5-(chloromethyl)pyrazolo[1,5-a]pyrimidine-3-carboxylate. Chloro-5-(chloromethyl)pyrazolo[1,5-a]pyrimidine-3-carboxylate (40 mg, 0.15 mmol) was dissolved in 1 ml of MeOH and NaOMe (17 mg, 0.31 mmol) was added at 0 °C. The reaction mixture was then stirred overnight at room temperature. The reaction was then evaporated, diluted with EtOAc and washed with water. The organic layer was dried over Mg_2_SO_4_, filtered, and concentrated under reduce pressure. The resulting residue was purified on a pre-packed Silica gel column on a Teledyne ISCO purification system to give desired product in 38% isolated yield. ^1^H NMR (600 MHz, CDCl_3_) δ 8.59 (s, 1H), 6.70 (s, 1H), 4.81 (s, 2H), 4.32 (s, 3H), 3.96 (s, 3H). ESI-MS (m/z): 256.0 [M+H]^+^. ^13^C NMR (151 MHz, CDCl_3_) δ 162.88, 162.35, 156.64, 148.65, 148.21, 102.72, 87.72, 57.70, 51.73, 46.45. ESI-MS (m/z): 256.0 [M+H]^+^

#### SW396272. Methyl 5-(1-chloroethyl)-7-hydroxypyrazolo[1,5-a]pyrimidine-3-carboxylate

Methyl 3-amino-1H-pyrazole-4-carboxylate (50 mg, 0.35 mmol) and ethyl 4-chloro-3-oxopentanoate (82 mg, 0.46 mmol) were stirred in 1.0 ml of AcOH at 80 °C for 4 hours. The reaction mixture was then cooled down to 0 °C and filtered to give desired product as a solid in 29 % isolated yield. ^1^H NMR (400 MHz, CDCl_3_) δ 9.88 (s, 1H), 8.10 (s, 1H), 5.97 (s, 1H), 4.99 (q, *J* = 6.9 Hz, 1H), 3.85 (s, 3H), 1.86 (d, *J* = 6.8 Hz, 3H). ^13^C NMR (151 MHz, CDCl_3_) δ 163.50, 155.67, 151.17, 143.63, 143.29, 97.97, 97.54, 52.88, 51.88, 24.17. ESI-MS (m/z): 256.0 [M+H]^+^.

#### SW396744. Methyl 5-cyano-7-hydroxypyrazolo[1,5-a]pyrimidine-3-carboxylate

To a stirred solution of methyl 5-formyl-7-hydroxypyrazolo[1,5-a]pyrimidine-3-carboxylate (20 mg, 90 ml) in 2 ml of THF is added 500 ml of ammonium hydroxide (30%> aq. solution), followed by iodine (30 mg, 0.12 mmol, 1.3 equiv.), which caused a dark brown color to appear. The resulting reaction mixture was stirred for 10 minutes and was analyzed by LCMS. LCMS showed that all starting material was consumed and more than 90% of desired product’s formation. The reaction was quenched by addition of NaHCO_3_ solution and Na_2_S_2_O_3_ solution. After stirring for 10 minutes at room temperature, the reaction was purified on a pre-packed reverse phase column on a Teledyne ISCO purification system to give desire product in 61 % isolated yield. ^1^H NMR (600 MHz, MeOD) δ 8.20 (s, 1H), 6.12 (s, 1H), 3.76 (s, 3H). ^13^C NMR (151 MHz, DMSO) δ 163.30, 157.09, 151.42, 145.34, 134.69, 118.73, 100.86, 99.51, 50.99. ESI-MS (m/z): 219.1 [M+H]^+^

#### SW461447. Methyl 5-ethynyl-7-hydroxypyrazolo[1,5-a]pyrimidine-3-carboxylate

To a solution of methyl 5-formyl-7-hydroxypyrazolo[1,5-a]pyrimidine-3-carboxylate (18 mg, 81µmol) and dimethyl (1-diazo-2-oxopropyl)phosphonate (31 mg, 0.16 µmol) in 1.0 ml MeOH was added K_2_CO_3_ (34 mg, 0.24 mmol). The reaction mixture was stirred at 20 °C for 2 hours before adding water and EtOAc. The combined organic layers were washed with brine, dried over anhydrous MgSO_4_, filtered and the filtrate was concentrated in vacuo. The crude product was purified on a pre-packed Silica gel column on a Teledyne ISCO purification system to give desired product in 50% isolated yield. ^1^H NMR (600 MHz, DMSO) δ 8.03 (s, 1H), 5.70 (s, 1H), 4.09 (s, 1H), 3.69 (s, 3H). ^13^C NMR (151 MHz, DMSO) δ 163.55, 157.39, 151.79, 144.77, 143.85, 99.98, 98.00, 84.66, 79.18, 50.60. ESI-MS (m/z): 218.1 [M+H]^+^.

#### SW461448. Methyl 5-chloro-7-hydroxypyrazolo[1,5-a]pyridine-3-carboxylate

^1^H NMR (400 MHz, DMSO) δ 8.03 (s, 1H), 5.56 (s, 1H), 3.70 (s, 3H). ESI-MS (m/z): 228.0 [M+1]^+^. Synthesized as described in published patent application CN114763360A (46).

#### SW398310. Methyl 7-hydroxy-5-(2-hydroxyethyl)pyrazolo[1,5-a]pyrimidine-3-carboxylate

Methyl 3-amino-1H-pyrazole-4-carboxylate was stirred in EtOAC at 60 °C until it was completely dissolved. The solution was then cooled down to room temperature and Pd/C (10%) was then added. The hydrogenation reaction was performed for 48 hours under 80-100 psi. ^1^H NMR (600 MHz, DMSO) δ 8.21 (s, 1H), 5.86 (s, 1H), 3.81 (s, 3H), 3.76 (t, *J* = 5.9 Hz, 2H), 2.86 (t, *J* = 5.9 Hz, 2H). ^13^C NMR (151 MHz, DMSO) δ 162.62, 155.98, 154.66, 144.02, 143.42, 98.97, 96.81, 59.87, 51.76, 35.41.). ESI-MS (m/z): 238.1.

#### SW398309. Methyl 5-(2-chloroethyl)-7-hydroxypyrazolo[1,5-a]pyrimidine-3-carboxylate

To the solution of methyl 7-hydroxy-5-(2-hydroxyethyl)pyrazolo[1,5-a]pyrimidine-3-carboxylate (20 mg, 84 μl) in 1 ml of AcCN was added triphenylphosphane (44 mg, 0.17 mmol) followed by CCl_4_ (39 mg, 0.25mmol). After stirring stirring at room temperature for 24 hours, the reaction mixture was diluted with EtOAc and washed with water. The organic layer was dried over Mg_2_SO_4_, filtered, and concentrated under reduce pressure. The resulting residue was purified on a pre-packed Silica gel column on a Teledyne ISCO purification system to give desire product in 20% isolated yield. ^1^H NMR (600 MHz, MeOD) δ 8.25 (s, 1H), 5.97 (s, 1H), 3.93 (t, *J* = 6.9 Hz, 2H), 3.89 (s, 3H), 3.16 (t, *J* = 7.0 Hz, 2H). ESI-MS (m/z): 256.0 [M+H]^+^

### Cryo-EM sample and grid preparation

6xHis-TEV-nsp12 was incubated with NCI-2 at a 1:2 nsp12:drug ratio for 15 minutes at room temperature. At the same time nsp7 and nsp8 were mixed together at a 1:1 nsp7:nsp8 ratio and incubated for 15 minutes at room temperature. nsp12/NCI-2 was then mixed with nsp7/8 at a 1:3 ratio of nsp12/NCI-2:nsp7/8 and incubated for 15 mins at room temperature to allow for the replication-transcription complex (RTC) to form. An annealed primer (5’-CGCGUAGCAUGCUACGUCAUUCUCCUAAGAAGCUG-3’) (Millipore Sigma) and template (5’-CUAUCCCCAUGUGAGCGGCUCAGCUUCUUAGGAGAAUGACGUAGCAUGCUACGCG-3’) (IDT) RNA scaffold was then added to the RTC at a 1:1.2 RTC:RNA ratio and incubated at for 15 min at room temperature. The assembled RTC was purified by size exclusion chromatography through a Superdex 200 10/300 increase column in 50 mM Tris pH 8.0, 100 mM NaCl, 1 mM MnCl2, and 1 mM DTT. The fractions containing the RTC were pooled, concentrated in an Amicon Ultra-15 spin concentrator with a 10 kDa MWCO, aliquoted, and stored at −80 °C.

The final RTC sample was diluted to 3.6 mg/mL and supplemented with 0.025% DDM detergent. Glow discharged Copper Quantifoil 1.2/1.3 mesh 300 grids were used to freeze 3.5 μL of sample at 100% relative humidity at 8°C using Vitrobot mk. IV.

### Cryo-EM data collection

Data collection was performed by Dr. Yang Li from UT Southwestern Structural Biology Laboratory (SBL). Data collection was performed on a Titan Krios microscope at the Cryo Electron Microscopy Facility (CEMF) at UT Southwestern Medical Center, with the post-column energy filter (Gatan) and a K3 direct detection camera (Gatan), using SerialEM (47). 6,714 movies were acquired at a pixel size of 0.42 A in super-resolution counting mode, with an accumulated total dose of ∼60 e^-^/Å^2^ over 60 frames. The defocus range of the images was set to be −1.1 to −2.4 μm.

### Image processing and 3D reconstruction

Data was processed using a mixed pipeline in Relion 4 (48) and CryoSPARC 4.2.1 (48). Bayesian polishing and 3D classifications were performed in Relion, while all the other steps were done in CryoSPARC. Movies were motion corrected using Relion implementation of MotionCor2 (49), with a downsampled pixel size of 0.84 Å. Aligned movies were imported into CryoSPARC for patch CTF estimation. Particles were picked using crYOLO(50), and CryoSPARC template picker. Templates were generated from PDB 7THM, and results of the initial 2D classifications were used for a second round of template picking. All three sets were classified separately, and then merged, removing the duplicates. Non-Uniform refinement was initiated with an Ab-initio model, followed by CTF refinement. Local refinement was performed with a mask encompassing the entire RTC. Aligned particles were imported into Relion for 3D classification with alignment using csparc2star scripts (51). Good classes were re-imported into CryoSPARC for alignment and exported into Relion for two rounds of Bayesian polishing. After additional 2D classification of the output particles and re-refinement, two rounds of alignment-free 3D classification were performed: Overall, and NiRAN-focused, with only best classes selected from both. Final maps were an output from CryoSPARC local refinement with a mask encompassing the entire RTC. Figures were prepared in CryoSPARC, Chimera (52), ChimeraX (53), and PyMOL (54).

It is worth noting that the final reconstruction is missing the globular portion of one of the nsp8 molecules. A focused classification of the region results in two conformations of the missing nsp8 – “up” position consistent with the published structures, and a “down” position. Since the NiRAN-bound inhibitor was the focus of the study, we did not exhaustively assess the nsp8 conformational dynamics.

### Model Building

The RNA duplex used in RTC reconstitution was based on the PDB 7UO4 (36), and thus the PDB file was used as the initial model. Model was rigid fit to the map in Chimera. Model was rebuilt/truncated in Coot(55) and optimised in ChimeraX/ISOLDE (56). Ligand and bond restraints were generated with AceDRG (57). SMILES string used was:

OC1=CC(C)=NC2=C(C=NN21)C(OC)=O

and was derived from the initial compound NCI-2 by removing the chlorine group

OC1=CC(CCl)=NC2=C(C=NN21)C(OC)=O

Solvent was built manually. Model refinement was performed against the sharpened map in Phenix (58) with initial model restrains, and ligand restraints from phenix elbow. Cys53-inhibitor S-C bond length was fixed based on the AceDRG output to 1.82 Å +/− 0.05 sigma in a manually prepared parameter file. Model validation was performed using MolProbity (59).

## Data availability

The atomic coordinates for NCI-2 bund to the RTC are deposited in the Protein Data Bank with entry code 9CGV. The maps were deposited in EMDB under accession EMD-45587. The data used to support the findings of the study are available from the corresponding author upon reasonable request.

## Supporting information

This article contains supporting information.

## Author Contributions

G.H. and V.S.T. conceived and designed study; G.H., H.N., B.P., and V.S.T. designed HTS assay; G.H., A.M., and H.N. prepared and performed HTS; G.H., A.M., H.N., B.P., J.A.R., and V.S.T. analyzed HTS data; M.A. and J.A.R. designed and synthesized NCI-2 derivatives; J.L.E. and J.W.S. designed, performed, and analyzed SARS-CoV-2 infection assays; K.A.S. performed mass spectrometry; G.H. and A.O. prepared samples for Cryo-EM; A.O. performed structural analysis; K.P. performed bioinformatics; G.H. performed all other biochemical and cell culture experiments; G.H., A.M., J.W.S., J.A.R., and V.S.T. analyzed data from biochemical and cell culture experiments; G.H. and V.S.T. wrote the manuscript with input from all authors.

## Funding and additional information

This work was funded by NIH Grants R01GM135189 (V.S.T.), a W. M. Keck Foundation Medical Research Grant (V.S.T., K.P., J.S.) a Welch Foundation Grants I-1911 (V.S.T.), I-1612 (J.M.R.) and AI158124 (J.W.S) and the Howard Hughes Medical Institute (HHMI, V.S.T). G.H. was supported by a Donald D. Brown Life Sciences Research Foundation Postdoctoral Fellowship. V.S.T. is a Michael L. Rosenberg Scholar in Medical Research, a CPRIT Scholar (RR150033), a Searle Scholar and an investigator of the HHMI. The Structural Biology Lab at UT Southwestern Medical Center is partially supported by grant RP220582 from the Cancer Prevention & Research Institute of Texas (CPRIT) and the Cryo-Electron Microscopy Facility at UT Southwestern Medical Center is partially supported by grant RP170644 from CPRIT. Drug screening and dose-response studies were carried out with an Echo 655, which was obtained by B.P. through a shared instrumentation grant (S10 OD026758-01).

## Acknowledgements

We thank members of the Tagliabracci laboratory for helpful discussions, Anju Sreelatha for SelO protein, Yang Li for structural help, Andrew Lemoff for intact mass analysis and Noelle Williams for support with LC-MS/MS.

