## Supporting information for "Covalent inhibition of the SARS-CoV-2 NiRAN domain via an active-site cysteine"

**Affiliations:**

*Correspondence to: Vincent S. Tagliabracci

#### Supplemental Figures

A

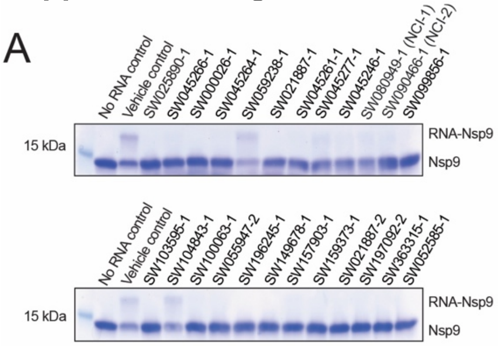


B

| UTSW ID | PAINS | DSF Peak Temp (°C) | ΔT (°C) |
|---|---|---|---|
| SW025890-1 | Yes | Undeterminable | |
| SW045266-1 | Yes | Undeterminable | |
| SW000026-1 | Yes | Undeterminable | |
| SW045264-1 | Yes | Undeterminable | |
| SW059238-1 | Not Pains | 43.25 | -0.5 |
| SW021887-1 | Yes | Undeterminable | |
| SW045261-1 | Yes | Undeterminable | |
| SW045277-1 | Yes | Undeterminable | |
| SW045246-1 | Yes | Undeterminable | |
| SW080949-1 | Not Pains | 44.75 | 1 |
| SW090466-1 | Yes | 45.25 | 1.5 |
| SW099856-1 | Not Pains | Undeterminable | |
| SW103595-1 | Yes | Undeterminable | |
| SW104843-1 | Not Pains | 43.25 | -0.5 |
| SW100063-1 | Not Pains | 43.25 | -0.5 |
| SW055947-2 | Yes | 39.75 | -4 |
| SW196245-1 | Not Pains | Undeterminable | |
| SW149678-1 | Yes | Undeterminable | |
| SW157903-1 | Yes | 45.25 | 1.5 |
| SW159373-1 | Yes | Undeterminable | |
| SW021887-2 | Yes | Undeterminable | |
| SW197092-2 | Not Pains | 35.25 | -8.5 |
| SW363315-1 | Not Pains | Undeterminable | |
| SW052585-1 | Yes | Undeterminable | |

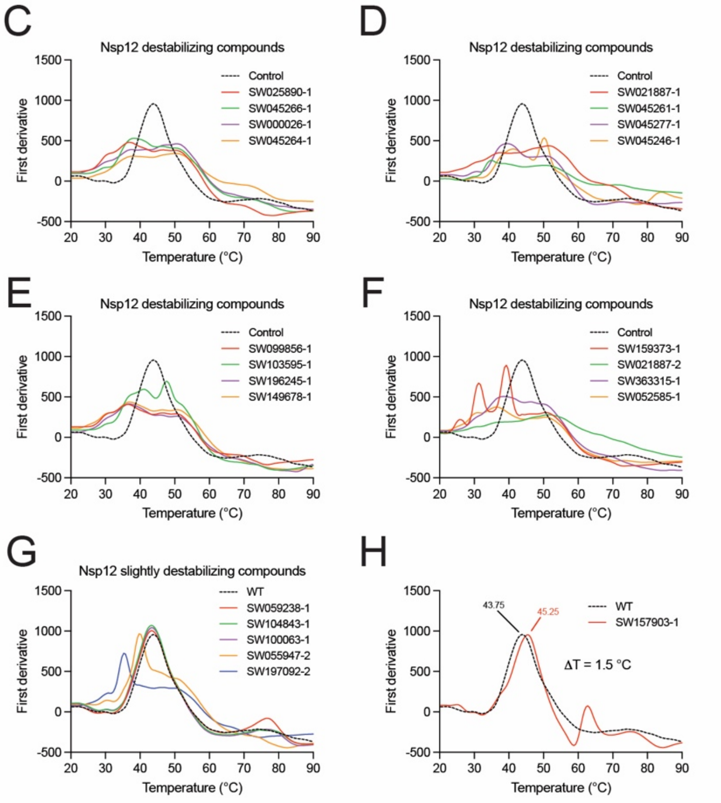


**Figure S1. Characterization of the small molecule identified in the HTS. A,** RNAylation activity of nsp12, (500 nM) following incubation with various small molecules (20 μM) identified in the HTS. Reactions were initiated with 100 μM of SARS-CoV-2 genomic leader sequence corresponding to the first 10 bases in the genome, and the reaction products were analyzed by SDS-PAGE and Coomassie staining. B, table depicting the ID, PAINS class, and thermal stability profile (peak temp. and change melting temperature; ΔT) of the small molecules identified from the HTS. C-H, differential scanning fluorimetry profiles of nsp12 (10 μM) in the presence of the small molecules identified from the HTS. Compounds were grouped into nsp12 destabilizing compounds (C-F), nsp12 slightly destabilizing compounds (G) and SW157903 (H), which stabilized nsp12; however, we did not follow up on due to its PAINS class.

| Modified Residue | Nsp12 Peptide | Spectral Count +NCI-1 | Spectral Count +NCI-2 |
|---|---|---|---|
| C22 | DAQSFLNRVCGVSAARLTPCGTGTST | 5 | 3 |
| C53 | DIYNDKVAGFAKFLKTNCCRFQEK | 6 | 0 |
| C53 | DKVAGFAKFLKTNCCRFQEK | 13 | 3 |
| C53 | DKVAGFAKFLKTNCCRFQEKDE | 2 | 0 |
| C53 | DKVAGFAKFLKTNCCRFQEKDEDDNLI | 0 | 7 |
| C93 | DCPAVAKH | 1 | 0 |
| C152 | DTLKEILVTYNCC | 2 | 0 |
| C395 | DKRTTCFSVAALTNNVAFQTVKPGNFNKDFY | 2 | 0 |
| C487 | DGGCINANQVIVNNL | 5 | 1 |
| C622 | DVENPHLMGWDYPKC | 2 | 0 |

**Figure S2. LC-MS/MS analysis of nsp12 bound to NCI-1 and NCI-2.** Table depicting the nsp12 peptides identified by LC-MS/MS to be covalently modified by NCI-1 (+192 Da) or NCI-2 (+205 Da) on cysteine residues. Modified cysteines are highlighted in red. Spectral counts reflect the number of MS2 peptide spectral matches as determined by Mascot software searches.

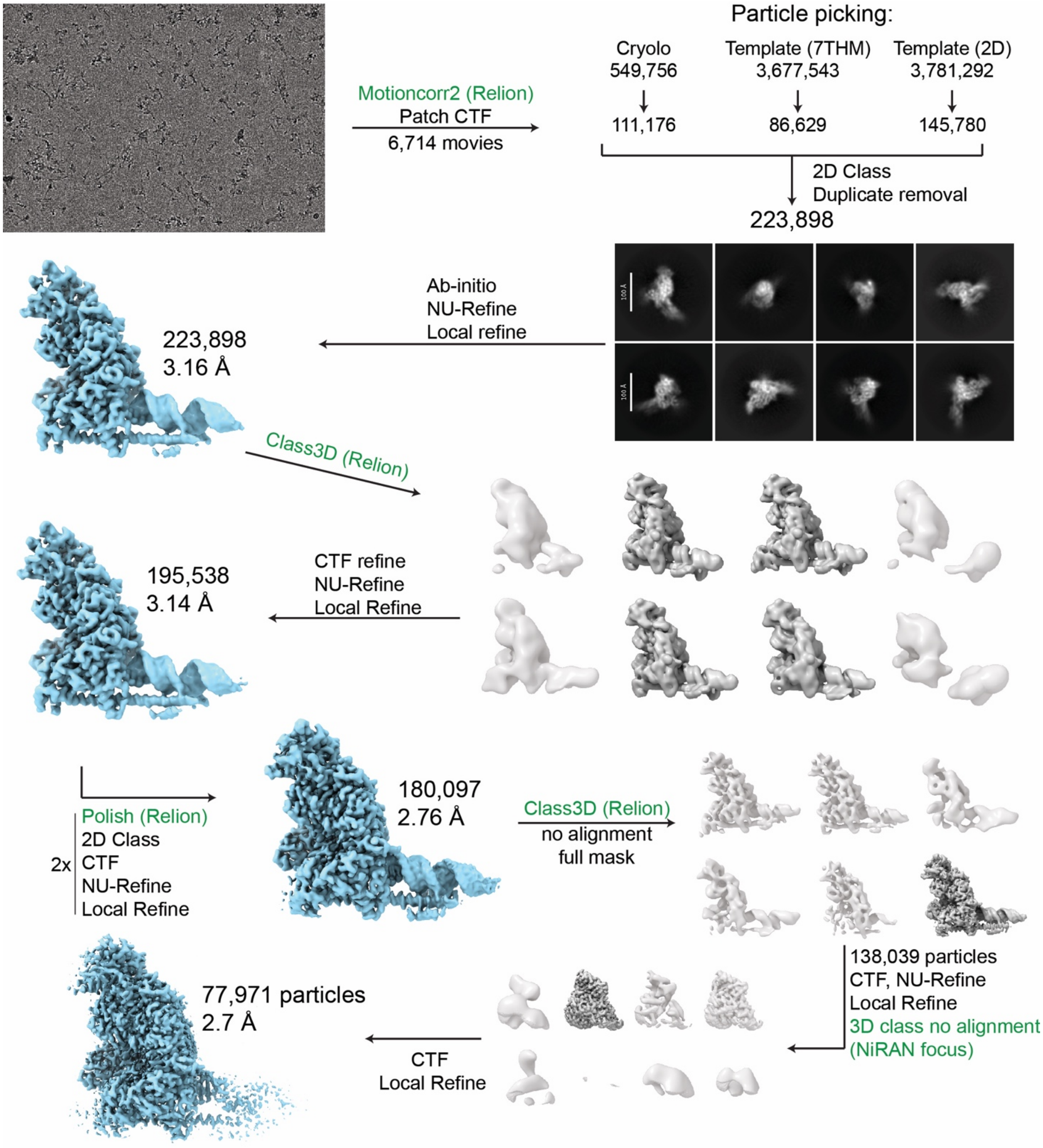


**Figure S3. Cryo-EM data processing scheme.** Data was processed in CryoSPARC 4.2.1. Steps indicated in green were performed in Relion 4. Particles were picked with CryoSPARC and crYOLO 1.9.3.

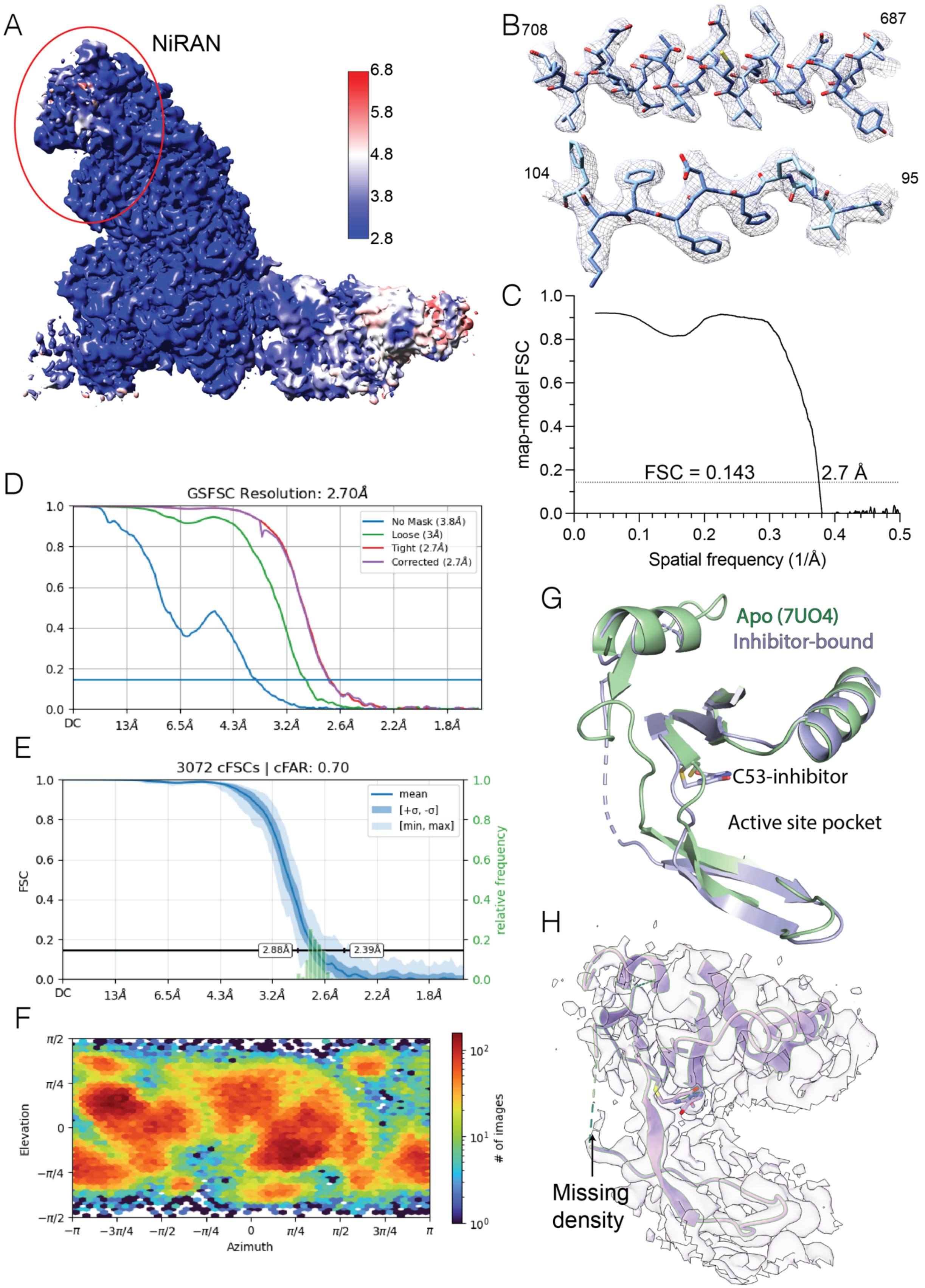
A
NiRAN
6.8
5.8
4.8
3.8
2.8
B
708
687
104
95
C
map-model FSC
FSC = 0.143
2.7 Å
Spatial frequency (1/Å)
D
GSFSC Resolution: 2.70Å
No Mask (3.8Å)
Loose (3Å)
Tight (2.7Å)
Corrected (2.7Å)
E
3072 cFSCs | cFAR: 0.70
mean
[+σ, -σ]
[min, max]
FSC
relative frequency
2.88Å
2.39Å
F
Elevation
Azimuth
### of images
G
Apo (7UO4)
Inhibitor-bound
C53-inhibitor
Active site pocket
H
Missing density

**Figure S4. Cryo-EM structure quality metrics.** A, local resolution estimation of the RTC bound to NCI-2. NiRAN domain is indicated. B, example coulombic map density and model fit within the polymerase domain (top) and the NiRAN domain (bottom). C, map-model FSC curve of the RTC bound to NCI-2. D, gold-standard FSC curves for the final refinement. E. The conical FSC Area Ratio plot. Reported Sampling Compensation Factor (SCF) was 0.902. F, particle angular distribution plot for the final refinement. G. Binding to the NCI-2 destabilizes a strand proximal to Cys53. The missing structure is presented as dashed lines on the left side of the panel. Light-blue cartoon of the inhibitor-bound structure is aligned with an apo structure in green; pdb: 7UO4 (36). H. A view of NiRAN structure with the missing density around the inhibitor binding site.
